# Mechanism of strand displacement DNA synthesis by the coordinated activities of human mitochondrial DNA polymerase and SSB

**DOI:** 10.1101/2022.07.19.500644

**Authors:** Plaza-G.A. Ismael, Kateryna M. Lemishko, Rodrigo Crespo, Thinh Q. Truong, Laurie S. Kaguni, Francisco J. Cao-García, Grzegorz L. Ciesielski, Borja Ibarra

**Author notes:** These authors contributed equally to these work (Plaza-G.A. Ismael, Kateryna M. Lemishko). Department of Physics, King’s College London, Strand Campus, London WC2R 2LS, UK.

## Abstract

Many replicative DNA polymerases couple DNA replication and unwinding activities to perform strand displacement DNA synthesis, a critical ability for DNA metabolism. Strand displacement is tightly regulated by partner proteins, such as single-stranded DNA (ssDNA) binding proteins (SSBs) by a poorly understood mechanism. Here, we use single-molecule optical tweezers and biochemical assays to elucidate the molecular mechanism of strand displacement DNA synthesis by the human mitochondrial DNA polymerase, Polγ, and its modulation by cognate and noncognate SSBs. We show that Polγ exhibits a robust DNA unwinding mechanism, which entails lowering the energy barrier for unwinding of the first base pair of the DNA fork junction, by ∼55%. However, the polymerase cannot prevent the reannealing of the parental strands efficiently, which limits by ∼30-fold its strand displacement activity. We demonstrate that SSBs stimulate the Polγ strand displacement activity through several mechanisms. SSB binding energy to ssDNA additionally increases the destabilization energy at the DNA junction, by ∼25%. Furthermore, SSB interactions with the displaced ssDNA reduce the DNA fork reannealing pressure on Polγ, in turn promoting the productive polymerization state by ∼3-fold. These stimulatory effects are enhanced by species-specific functional interactions and have significant implications in the replication of the human mitochondrial DNA.

## INTRODUCTION

Replication of mitochondrial DNA (mtDNA) is carried out by specialized replication machinery (replisome). *In vitro*, three components of the human mitochondrial replisome are sufficient for the synthesis of the genome-long DNA over primed templates: the DNA polymerase holoenzyme, Polγ, the mitochondrial single-stranded DNA-binding protein (mtSSB) and the mitochondrial DNA helicase (TWINKLE) (1-3). Defects in the operation of any of these proteins have been linked to human diseases and ageing (4-7). Polγ is a heterotrimeric holoenzyme that consists of a catalytic subunit (PolγA) and a dimeric accessory subunit (PolγB), Figure 1A (8,9). The catalytic subunit exhibits a finely tuned balance of polymerase (*pol*) and 3’-5’ exonuclease (*exo*) activities that ensures efficiency as well as fidelity of DNA synthesis (9-12). As in the case of other replicative DNA polymerases (DNApols) (13-15), Polγ also exhibits an intrinsic strand displacement DNA synthesis activity, which entails coupling of the DNA synthesis and unwinding activities (8,16).

**Figure 1:**
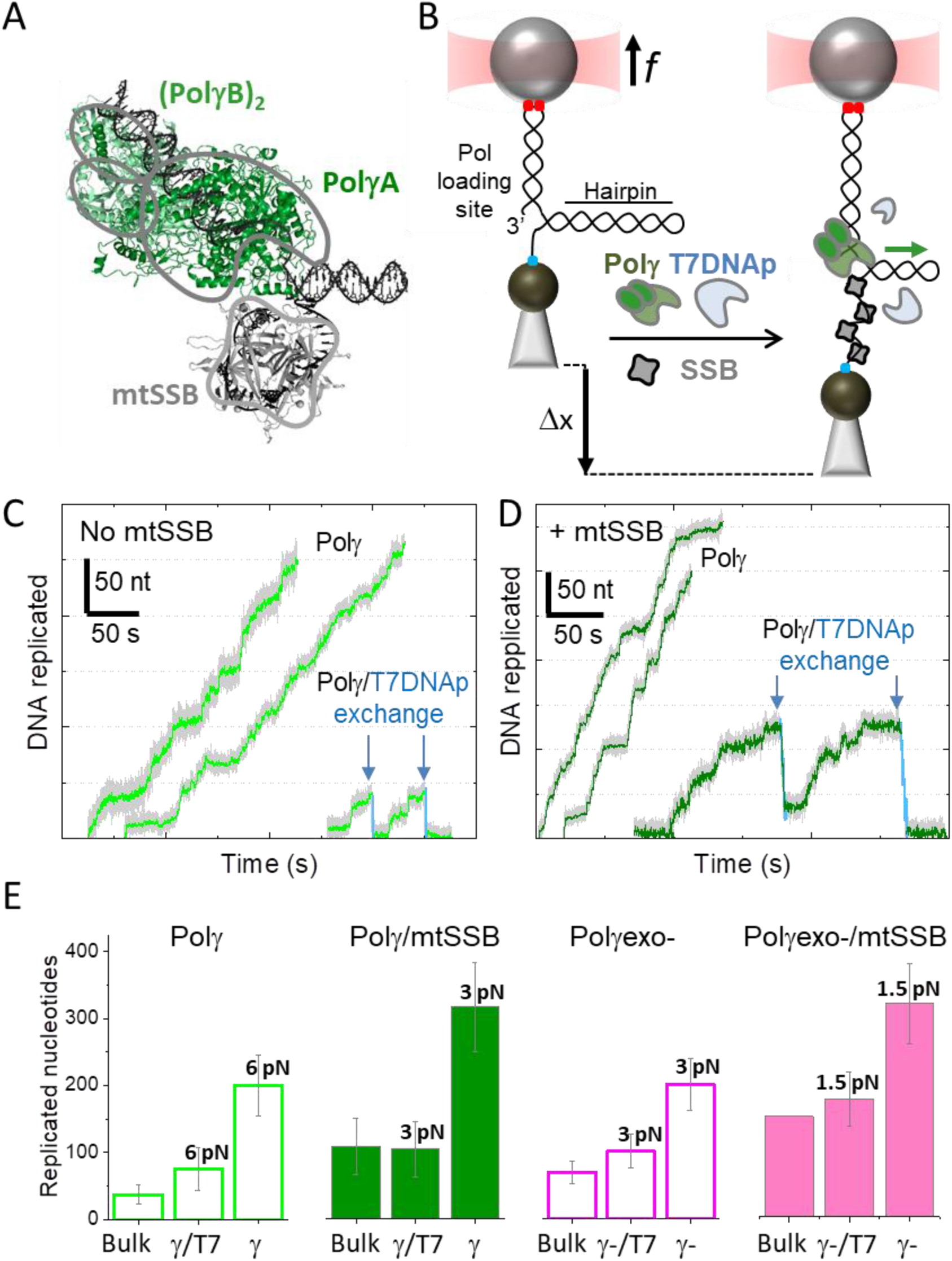
Polγ and Polγexo- exchange with T7DNAp in solution. **A)** Schematic of Polγ (10.2210/pdb3IKM/pdb) and mtSSB (10.2210/pdb3ULL/pdb) at the DNA fork. Polγ holoenzyme is composed by the catalytic subunit, PolγA (dark green), and a dimer of the accessory subunit, PolγB, (light green). mtSSB (grey) binds the displaced ssDNA as a tetramer. **B)** In the optical tweezers, a single DNA hairpin (559 bp) is tethered between two functionalized beads and held at constant tension (*f*). One strand of the hairpin is connected to the bead in the optical trap (red cone) through a dsDNA handle (via digoxigenin-antibody connections, red dots). The other strand is attached to a bead on a micropipette by biotin-streptavidin linkages (blue dot). The dsDNA handle includes a 3’ end for polymerase loading, and the 5’ end of the hairpin includes a poly-(dT)_30_ site for SSB binding. Strand displacement DNA synthesis by Polγ (or Polγexo-) increases the end-to-end extension of the hairpin (Δx). Green arrow indicates the 5’ to 3’ direction of DNA polymerase translocation along the hairpin. Experiments were performed without and with several concentrations of cognate and noncognate SSBs (grey) and/or competitor T7DNAp (light blue) in solution. **C)** Representative experimental traces of Polγ (2 nM) without (left) and with (right) competing T7DNAp (1 nM) in solution. **D)** Traces of Polγ (2 nM) activity in the presence of mtSSB (50 nM) without (left) and with (right) competing T7DNAp (1 nM) in solution. Exchange of Polγ by T7DNAp was monitored as *exo* events (blue) not observed in the absence of T7DNAp (*f*= 6-7 pN). **D)** Comparison of the maximum number of replicated nucleotides detected in single turn over bulk experiments (Figure S2) with the average number of replicated nucleotides measured in optical tweezers assays at the lowest detection tensions in the absence and presence of mtSSB. For each panel, bulk data (Bulk), Polγ (γ) and Polγexo- (γ-). Single-molecule data in the presence of T7DNAp is labelled as γ/T7 or γ-/T7. Error bars show standard errors.

Strand displacement DNA synthesis by Polγ is relevant for replication through stable secondary structures (17-19), maintenance of the D-loop DNA structure at the origin of replication of the heavy strand (20), and removing RNA/DNA primers in coordination with primer processing factors (21-23). Similarly to other DNApols (13,15,24), the efficiency of strand displacement activity of Polγ is limited to a few nucleotides (16,22,25-27). Biochemical studies have shown that at a nick Polγ is prone to enter the idling state, characterized by repeated addition and excision of a nucleotide, which is driven by the intramolecular and reversible transfer of the primer between the *pol* and the *exo* active sites (11,16). According to previous single-molecule manipulation studies on other DNApols, the regression or reannealing pressure of the DNA fork during strand displacement would inhibit polymerization-driven forward motion and promote the partition of the primer to the *exo* domain (24,28). The *exo* activity would then limit the strand displacement levels. In fact, mutations that impede *exo* activity of Polγ stimulate its strand displacement activity (16,29). This enhanced strand displacement activity has detrimental physiological consequences. For example, during primer removal processes, excessive strand displacement hinders the coordination of the holoenzyme activity with that of the other primer processing factors (nucleases, helicases and/or mtSSBs), which has been linked with the formation of persistent unligatable flaps and double stranded breaks during mtDNA replication found in age-related pathologies *in vivo* (25,27,30).

Furthermore, recent studies have shown that DNA synthesis by replicative DNApols of phages T4 and T7 drives the fork unwinding during leading strand DNA replication (31,32). This activity is enhanced by other protein partners at the fork such as, the replicative helicase and SSBs (32,33). Considering the likely bacteriophage ancestry of the mtDNA replisome (34-36), it is tempting to speculate that the intrinsic DNA unwinding activity of Polγ contributes to the replication of the heavy strand of mtDNA significantly, while mtSSB and the helicase Twinkle would modulate the propensity of Polγ to idle at the fork junction in order to promote efficient DNA synthesis without compromising fidelity.

Despite the putative relevance of the Polγ ability to couple DNA synthesis and unwinding activities for mtDNA replication, little is known about the kinetics and mechanistic aspects of this reaction and it’s modulation by proteins partners, such as the mtSSB, to define the context dependent role of the holoenzyme. Human mtSSB is essential for mtDNA synthesis *in vitro* and *in vivo* (3,18,37). It binds ssDNA in a sequence independent manner (38,39) and forms the central nucleo-protein complex substrate upon which the mitochondrial polymerase must act (18,40). Although physical interactions between Polγ and mtSSB have not been observed (41), mtSSB can stimulate the primer-extension activity of Polγ (17,42). In other DNA replication systems SSBs have been shown to stimulate the strand displacement activity of DNApols (13,14,33,43,44). However, the effect of mtSSB on the strand displacement activity of Polγ remains largely unexplored.

Here, we present single-molecule manipulation optical tweezers assays, supported by ensemble biochemical experiments, to quantify the intrinsic strand displacement mechanism of Polγ and its modulation by mtSSB. We compare the real-time kinetics of the wild-type holoenzyme Polγ with that of an *exo* deficient variant, D_198_AE_200_A (45) (Polγexo-), in order to determine the contribution of the *exo* reaction on the strand displacement activity (without interference of idling events). In addition, to determine the putative role of species-specific polymerase-SSB interactions, we studied and compared the effects of several concentrations of cognate mtSSB and noncognate Escherichia coli (EcoSSB) and phage T7 (gp2.5) SSBs on the strand displacement activities of both holoenzyme variants. EcoSSB shares significant sequence and structural homology to mtSSB (46,47). Both proteins bind preformed ssDNA as tetramers with similar affinities (K_d_ ∼2 nM) and footprints (number of nucleotides wrapped per tetramer) (38,39,48-52). In contrast, the multifunctional gp2.5 is organized as a dimer, shows smaller ssDNA binding footprint and lower affinity for ssDNA (*K*_*D*_ ∼0.8 μM) than mtSSB and EcoSSB (53,54). EcoSSB and gp2.5 present intrinsically disordered acidic C-terminal tails that mediate interactions with other proteins including some of their respective replisomes (50,55). A comparable C-terminal tail is absent on mtSSB. Experiments were performed in the absence and presence of the phage T7 DNA polymerase (T7DNAp) in solution. Inspired by previous bulk and single-molecule studies in the field (56,57), we used the marked differences between T7DNAp and Polγ (and Polγexo-) strand displacement and *exo* rates as reporters to show that Polγ can exchange with competing DNApols in solution during active DNA synthesis and the exchange reaction is not rate-limiting.

Overall, our results show that Polγ presents a robust DNA displacement mechanism that is limited by reannealing of the parental strands, which shifts its activity equilibrium towards the *exo* state. We demonstrate that SSBs use several mechanisms to stimulate the strand displacement DNA synthesis by Polγ; i.e., mtSSB binding to the displaced ssDNA imposes additional destabilization energy on the DNA junction and reduces the DNA fork regression pressure on the holoenzyme. Interestingly, species-specific functional interactions enhance these stimulatory effects and thus may be critical under suboptimal mtSSB concentrations or stress conditions. Our measurements shed new light on the mechanism by which accessory proteins, such as SSBs, can enhance strand displacement DNA synthesis.

## MATERIALS AND METHODS

### Proteins and DNA constructs

Recombinant catalytic subunits (Polγ A) of wild-type and mutant (D_198_AE_200_A) Polγexo- variants were prepared from S*f9* cells (58). The accessory subunit of the holoenzyme (Polγ B) was prepared from bacterial cells (58). The catalytic and accessory subunits were combined in a 1:1.5 molar ratio to reconstitute the holoenzyme. Recombinant mtSSB was prepared from bacterial cells as described previously (42). Recombinant EcoSSB and Sequenase© were purchased from Thermofisher. T7DNAp was purchased from NEB. Recombinant gp2.5 was purchased from LSBio and Monserte Biotechnology.

The hairpin construct was synthesized as described previously (28). The construct consists of a 2,686 base pairs (bp) DNA ‘handle’ (pUC19 vector, Novagen) labeled with digoxigenin at one end, a 5’ (dT)_35_ end functionalized with biotin, and a 556 bp stem capped by a (dT)_4_ loop. The final hairpin construct contains a unique 3’ end loading site for the DNA polymerase (Figure 1B). The hairpin stem sequence is described in (28) and contains a 75% AT sequence. Considering the free energy formations of AT and GC bp as Δ*G*_*AT*_∼1.5 *k*_*B*_*T* and Δ*G*_*GC*_∼2.9 *k*_*B*_*T* (under ionic conditions similar to those used in this work) the average free energy of bp formation was Δ*G*_*bp*_∼1.8 *k*_*B*_*T* (59). For primer extension experiments, we used a gapped DNA template, consisting of ∼900 nucleotides of single-stranded DNA (ssDNA) flanked by ∼3550 bp dsDNA handles labeled with biotin and digoxigenin, as described in (60) (Figure S1).

### Optical tweezers experiments

We used a miniaturized counter propagating dual-beam optical tweezers instrument (61) to manipulate individual DNA hairpins tethered between a streptavidin-coated bead (2.1 μm, Kisker Biotech) immobilized on top of a micropipette and an anti-digoxigenin-coated bead (3.0 μm diameter, Kisker Biotech) held in the optical trap, Figure 1B (62). Proteins were introduced inside the flow cell after dilution in the replication buffer containing 50 mM Tris pH 8.5, 30 mM KCl, 10 mM DTT, 4 mM MgCl_2_, 0.2 mg/ml BSA and the four dNTPs (50 μM). Unless otherwise indicated, Polγ, Polγexo-, T7DNAp and Sequenase© were diluted to 2 nM. Polymerase exchange experiments were performed in a mixture of 2 nM Polγ or Polγexo- and 1 or 2 nM T7DNAp containing the indicated amounts of SSBs. Primer extension activities of Polγexo- diluted to 2 nM in the replication buffer were recorded on the gapped DNA construct, as described elsewhere (17,60), Figure S1. In all cases, data was monitored at 500 Hz at 22 ± 1 °C using a feedback loop to maintain a constant force or constant mechanical tension on the DNA. Force ranged explored were 1-11 picoNewtons (pN) in strand displacement assays and 1-16 pN in primer extension assays. The trap stiffness calibrated for 3.0 µm beads was *k* = 0.135±0.0043 pN nm^-1^

### Bulk biochemical experiments

Polγ strand displacement processivity and fork residence time assays in bulk were carried out in the replication buffer on a forked DNA substrate resembling the organization of the hairpin used in optical tweezers (Supplementary Methods and Figure S2).

### Data analysis

#### Processivity

The number of replicated nucleotides (processivity) in individual strand displacement assays was obtained by dividing the increase of the tether extension (Δx in Figure 1B) by the change in extension at a given tension accompanying during each catalytic step the generation of one new bp and one SSB-free or SSB-bound single-stranded nucleotide. The number of nucleotides incorporated in primer extension assays were obtained by dividing the change in tether extension by the change in extension due to the conversion of one single-stranded nucleotide into its double-stranded counterpart at a given tension (48,63). The extension of the dsDNA was approximated with the worm-like chain model for polymer elasticity with a persistent length of *P*= 53 nm and stretch modulus *S* = 1200 pN/ nm (64). The average extensions per nucleotide as a function of tension of free-ssDNA and ssDNA bound to mtSSB-, EcoSSB- or gp2.5-, under experimental conditions identical to those used in this work, were reported by us previously (17,51,65,66).

#### Average replication rates with and without pauses

The average replication rate at each tension (*V*_*mean*_(*f*)) was determined by a line fit to the traces showing the number of replicated nucleotides versus time. The final average rate at each tension was obtained by averaging over all of the traces taken within similar tension values (± 0.5 pN). Average replication rate without pauses at each tension (pause-free velocity, *V*(*f*)) was determined with an algorithm that computes the instantaneous velocities of the trajectory, averaging the position of the holoenzyme along the DNA over sliding time windows, as described previously (17). Tension dependent pause-free velocities were fitted to the strand displacement model described in Supporting Information (SI) and (67).

#### Average residence time at the pause state per nucleotide

The intrinsic flexibility of ssDNA together with the slow average strand displacement rates of the polymerases used in this work hindered the accurate identification of pause events (17). Nevertheless, identification of pause-free velocities allowed us to calculate the average residence time at the pause state per nucleotide at each tension, *T*_*p*_(*f*), as the difference between the total residence time per nucleotide (*T*_*t*_(*f*)= 1/ *V*_*mean*_(*f*)) and the average residence time in the active state (*T*_*a*_(*f*)= 1/ *V*(*f*)). The particular tension dependencies of *T*_*p*_(*f*) of each polymerase under study were fitted with *Eq*. 1. *T*_*p*_(*f*) can also be expressed in terms of *moving probability* (*MP*(f)), or the probability of finding the holoenzyme moving through the DNA hairpin as a function of tension. *MP*(f) was calculated as the ratio between the average replication rates with and without pauses at each tension, *MP*(*f*)= *V*_*mean*_(*f*)/ *V*(*f*), which is equivalent to *MP*(*f*)= *T*_*a*_(*f*)/ *T*_*mean*_(*f*) (see SI and (67)).

#### Maximum average processivities in the absence of tension

were estimated by multiplying the average residence time each holoenzyme spends at the fork (calculated from single turn over bulk experiments, 1/k_off_, Table 1) by their corresponding average velocities in the absence of tension, *V*_*mean*_(0). The latter value was assessed from values of *V*(0) and *T*_*p*_(0) obtained from the fits to the data with the 2-state and strand displacement models, respectively, *V*_*mean*_(0)= 1/(*T*_*a*_(0)+*T*_*p*_(0).

**Table 1.**
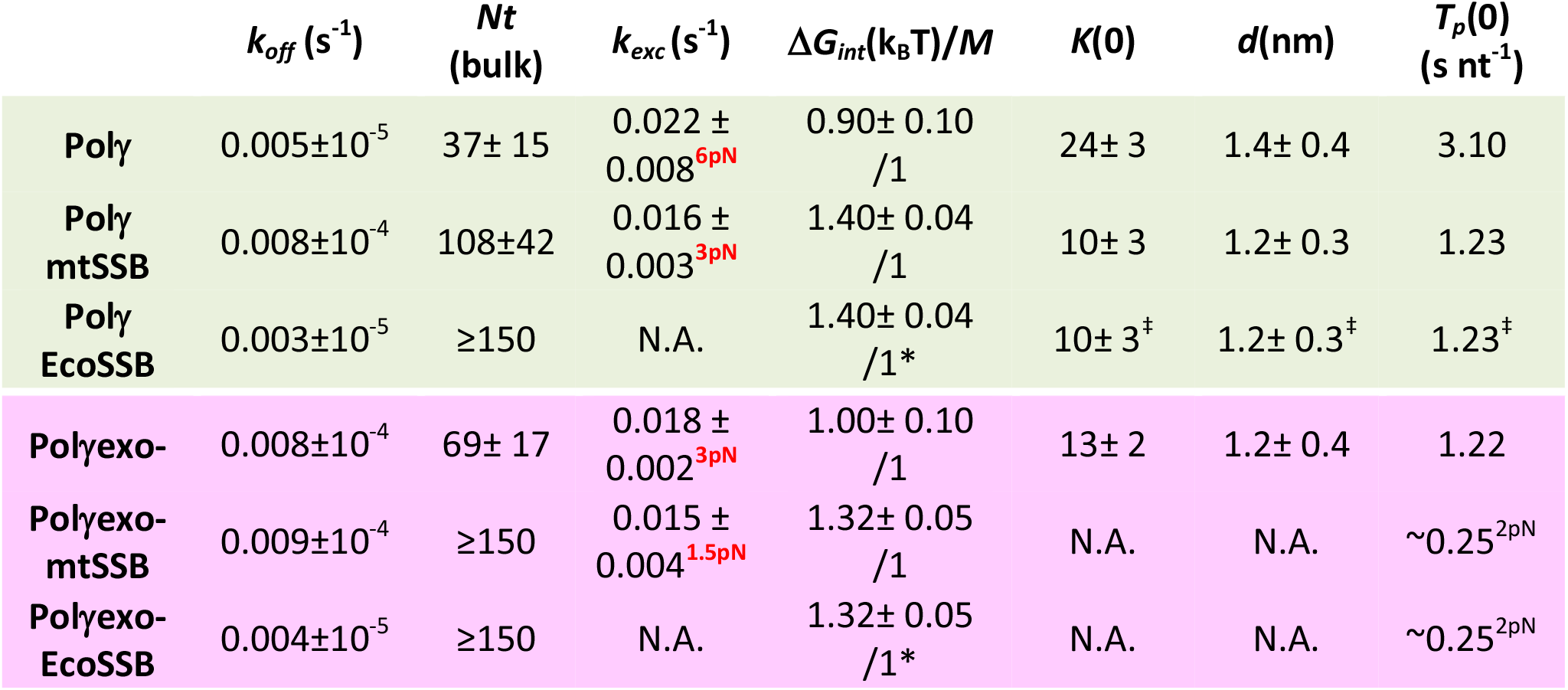
*k*_*off*,_ apparent detachment rates, and *Nt*, maximun number of replicated nucleotides determined in strand displacement bulk assays. *k*_*exc*_ shows the exchange rates of Polγ with T7DNAp (1:0.5 molar ratio) under mechanical tension (in red). Δ*G*_*int*_/*M*, minimum values of the free-parameters yielded by least squared error fits of strand displacement model to Polγ and Polγexo- pause-free velocity data. (*) Values from fits of the strand displacement model to Polγ and Polγexo- pause-free velocity data in the presence of mtSSB. *K*(0) and *d*, free-parameters of *Eq*.1 upon fitting experimental data with the least mean squared error.(^‡^) Values from fits of *Eq*.1 to Polγ residence time in pause state data in the presence of mtSSB. *T*_*p*_(0), average residence times at pause state in the absence of tension (0pN).(^2pN^) average residence times at pause state at ∼2 pN. In all cases, errors show standard errors. For Polγ, values correspond to conditions with 50 nM mtSSB or EcoSSB. For Polγexo- values correspond to conditions with 5 nM mtSSB and 50 nM EcoSSB.

## RESULTS

### Single-molecule strand displacement DNA synthesis assays

We used optical tweezers to follow the strand displacement DNA synthesis activity of Polγ holoenzyme and its *exo* deficient variant, Polγexo- (D_198_AE_200_A), on individual DNA hairpins, in real-time. The hairpin (559 bp) was flanked by 2.6-kb dsDNA and 30-nt ssDNA handles and tethered between two polystyrene beads, one held in an optical trap and the other fixed by suction on top of a micropipette, Figure 1B (Methods). The 3’ end of the 2.6-kb dsDNA handle was used as the polymerase holoenzyme loading site. Strand displacement activities were monitored at constant mechanical tension below 12 pN. Under these conditions, the end-to-end distance between the beads increases gradually as the mitochondrial holoenzyme replicates through the hairpin stem converting each DNA bp unwound to 1 dsDNA bp and 1 ssDNA nucleotide (Figures 1B-D and Methods). Experiments were carried out in the absence or presence of competing T7DNAp and several concentrations of mtSSB, EcoSSB, or gp2.5 diluted in the reaction buffer (Methods). We note that the hairpin was stably closed below 12 pN and, binding of either SSB to the ssDNA handle did not cause detectable DNA unwinding in the absence of the holoenzyme. For proper interpretation of the effect of mechanical tension on strand displacement activities, we performed independent measurements of the effect of tension on the primer extension replication kinetics of Polγ (17) and Polγexo- (Figure S1) holoenzymes.

### Polγ and Polγexo- exchange with competing polymerases in solution during strand displacement DNA synthesis

Single-molecule strand displacement activities were detected upon application of tension favoring fork destabilization, ∼3 and ∼6 pN for Polγexo- and Polγ, respectively. Notably, at the lowest tensions that allowed detection of activities, both holoenzymes already replicated hundreds of nucleotides (∼200 nt, Figures 1E, S3 and S4), which contrasted with the maximum processivity measured for each holoenzyme in bulk single turn over assays, ∼37 nt for Polγ and ∼69 nt for Polγexo- (Figures 1E, S2 and Table 1). These results suggested that the replication events measured in the optical tweezers may correspond to the consecutive action of several molecules that may exchange at the fork.

To determine if Polγ and Polγexo- could exchange with polymerase competitors in solution during active DNA synthesis, we measured (and compared) the real-time strand displacement kinetics of each holoenzyme at increasing tension in the absence and presence of competing T7DNAp (Methods). T7DNAp shares high level of structure and sequence similarity to the catalytic subunit of the mitochondrial Polγ holoenzyme (PolγA, Figure 1A) and almost identical DNA binding affinity (∼3 nM) (68,69). However, at tension below 8-10 pN, a stably closed hairpin prevents significant strand displacement DNA synthesis by T7DNAp and promotes its potent *exo* activity, which was detected as a fast and continuous decrease in the end-to-end distance of the DNA as the hairpin reanneals, Figures 1C, 1D and S3 (24). These characteristic *exo* events, which were not observed for the mitochondrial holoenzyme in the absence of T7DNAp at any tension, were used as reporters to identify the exchange of Polγ or Polγexo- with competing T7DNAp in solution during strand displacement DNA synthesis in the absence or presence of mtSSB (Figures 1C, 1D and S3). We note that at tension above 8-10 pN, polymerase exchange was monitored as a sudden increase in the replication rate due to the ∼4-times faster average replication rate of the phage DNA polymerase with respect to that of the mitochondrial holoenzyme at these tensions (Figure S3C).

At a molar ratio of 1 Polγ : 0.5 T7DNAp holoenzymes in solution, ∼80% of Polγ and Polγexo- traces showed long *exo* events indicating that the mitochondrial holoenzyme at the DNA fork can exchange with T7DNAp in solution at all tensions (Figures 1C, 1D and S3). Under these conditions, the average processivities of Polγ and Polγexo- (in the absence and presence of mtSSB) were 2-3 times shorter than those in the absence of T7DNAp and approached, at the lowest detection tensions, those measured in biochemical strand displacement replication assays under single turn over conditions (Figure 1E). These results suggest that, in the presence of competing T7DNAp in solution, strand displacement traces by Polγ or Polγexo- (in the absence and presence of mtSSB) would correspond to the activity of ∼1 holoenzyme, while in the absence of T7DNAp replication traces would correspond to the consecutive activity of several mitochondrial holoenzymes (2-3) that may exchange at the fork. Data showed in Figures 2, 3 and 4 were obtained in the presence of T7DNAp in solution (2 nM Polγ: 1 nM T7DNAp), results in the absence of T7DNAp are shown in Figure S4.

**Figure 2:**
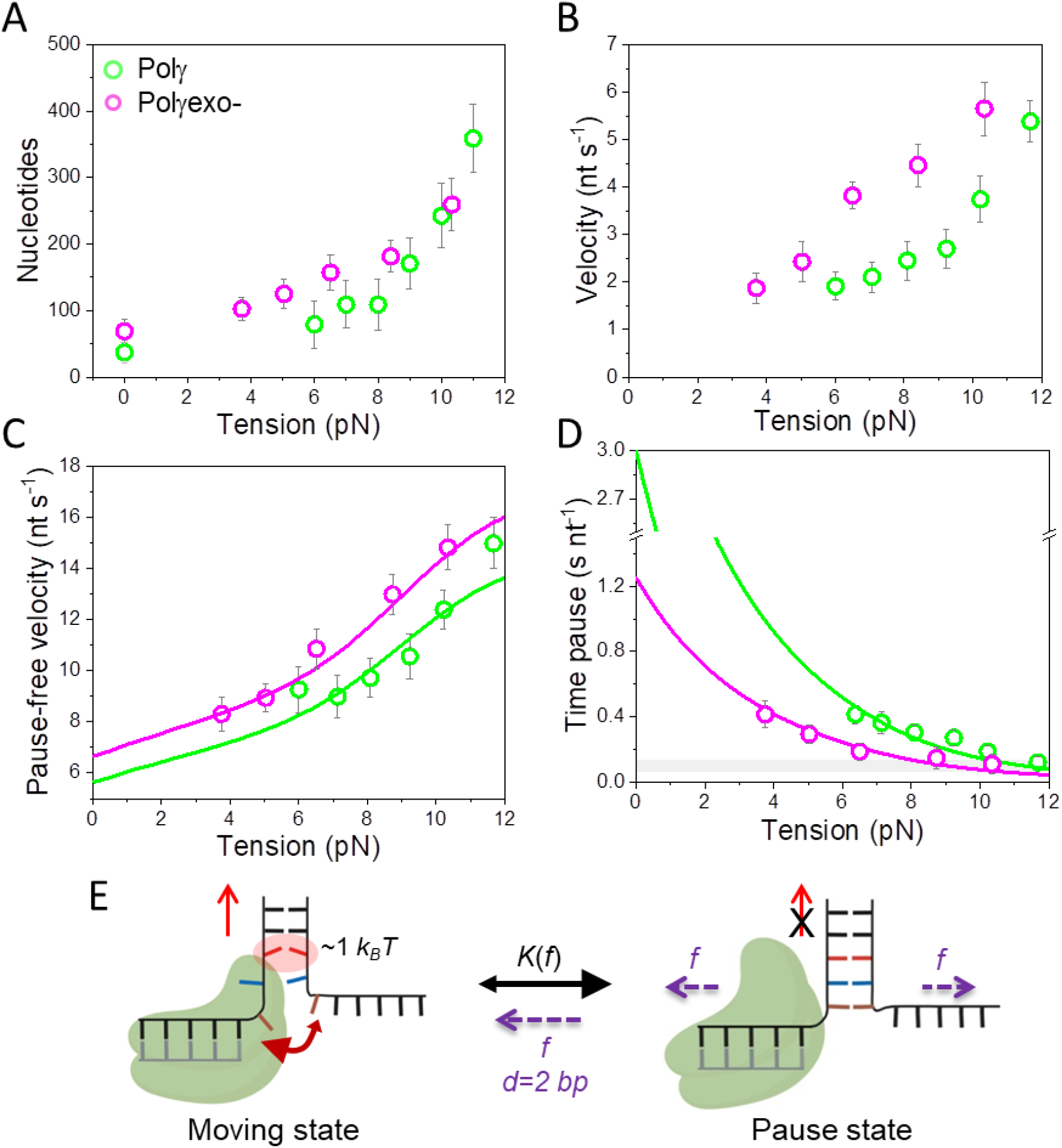
Effect of tension on Polγ and Polγexo- strand displacement kinetics. For all plots: Polγ green symbols (N=80), Polγexo- magenta symbols (N=71). Error bars show standard errors. For both holoenzymes **A**) the number of replicated nucleotides, and **B**) the average velocities (nt s^-1^) increase with tension (pN) continuously. Values at 0 pN correspond to bulk measurements. **C)** For both holoenzymes pause-free velocities (nt s^-1^) increased with tension continuously towards values measured during primer extension (Figure S1C). Green and magenta lines are the fits of the strand displacement model (SI) to Polγ and Polγexo- data, respectively. **D)** Tension dependencies of the average residence times at the pause state per nucleotide (*T*_*p*_(*f*), s nt^-1^). Green and magenta lines are the fits to Polγ and Polγexo- data, respectively, with a two state model (*Eq*. 1). Grey box show average *T*_*p*_(*f*) values measured under primer extension conditions in the absence of SSB, Figure S1D and (16). **E)** Diagram illustrating the two state model in which the holoenzyme alternates between moving and pause or non-productive states during the strand displacement reaction. In the moving state, two template nucleotides are bound to the *pol* site and the holoenzyme advances through the dsDNA destabilizing partially the first base pair of the junction with interaction energy of ΔG_int_ ∼1 *k*_*B*_*T* per dNTP incorporation step. In the absence of tension, the regression pressure of the dsDNA fork outcompetes the holoenzyme for the template, which shifts the equilibrium towards the pause or non-productive state strongly (*K*(0), Table 1) and reduces the probability of finding Polγ and Polγexo- in the moving state to ∼4 and 10%, respectively (SI). Application of tension (*f*) to the hairpin decreases the rewinding and/or favors the unwinding of first ∼2 bp of the fork (*d*), which shifts the equilibrium towards the moving state.

**Figure 3:**
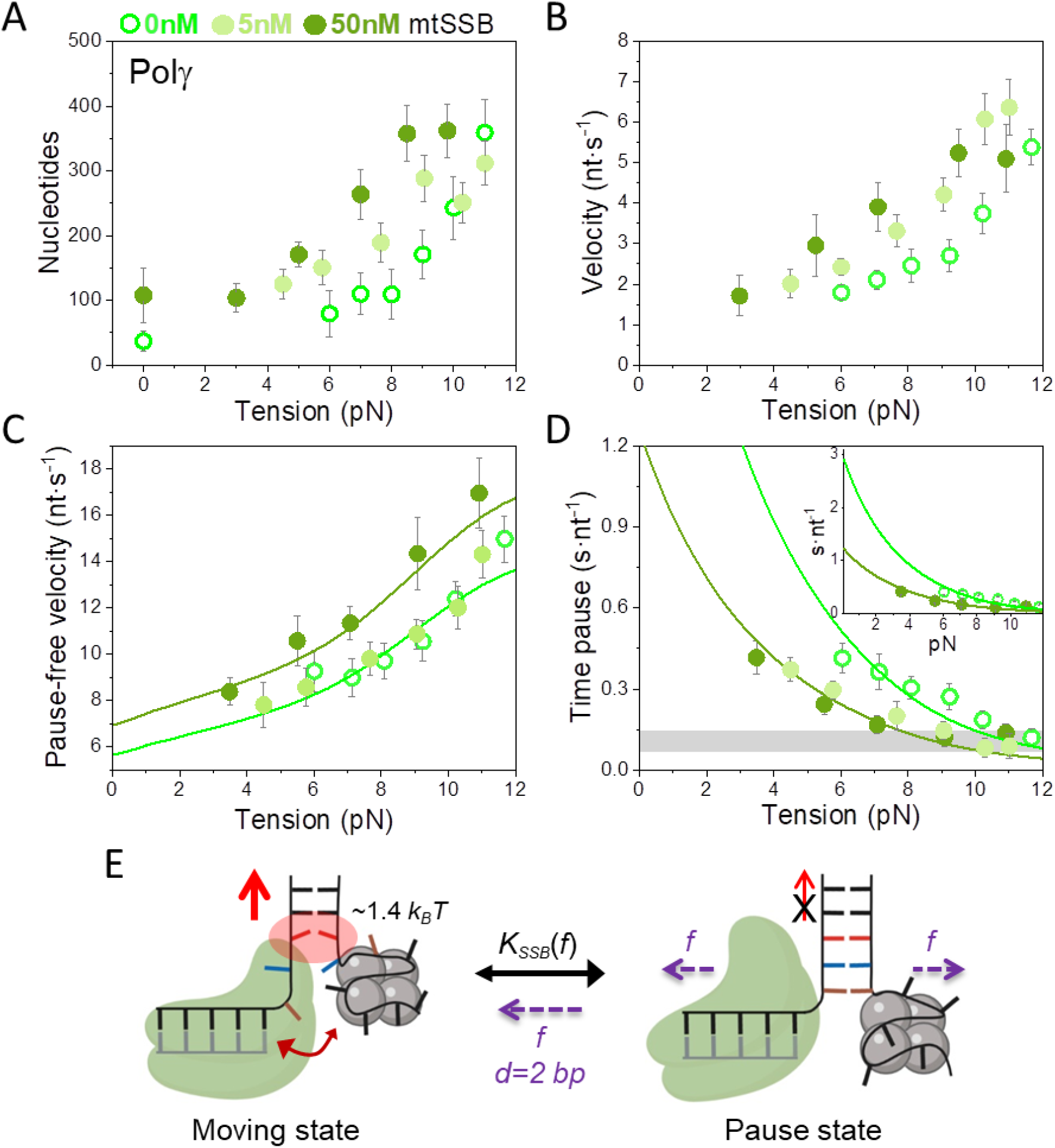
Effect of mtSSB on the tension dependent strand displacement kinetics of Polγ. Stimulatory effects of 5nM (N=20) and 50 nM (N=40) mtSSB on **A**) the average number of replicated nucleotides and **B**) average strand displacement rates (nt s^-1^) of Polγ as a function of tension (pN). **C**) mtSSB stimulates the pause-free velocity of Polγ at all tensions at 50 nM but not at 5 nM concentration. Light and dark green lines correspond to the fits of the strand displacement model to data in the absence and presence of mtSSB (50 nM), respectively. **D**) 5 and 50 nM concentrations of mtSSB decreased average residence time at pause state per nucleotide (*T*_*p*_(*f*), s nt^-1^) of Polγ at all tensions. Grey box shows average *T*_*p*_(*f*) values obtained under primer extension conditions in the absence of SSB (16). Light and dark green lines are the fits of two-state model (*Eq*.1) to data in the absence and presence of mtSSB (50 nM), respectively. mtSSB binding to the displaced strand decreases ∼2-3 times the average residence time of Polγ at a pause or non-productive state. **E**) Diagram illustrating the two-state model in the presence of mtSSB in which Polγ alternates between moving and pause or non-productive states. In the moving state, two template nucleotides are bound at the *pol* active site and the holoenzyme-mtSSB complex destabilizes partially the first base pair of the DNA hairpin with interaction energy a ∼40% higher than in the absence of SSB (Δ*G*_*int*_ ∼1.4 *k*_*B*_*T* per dNTP incorporated). In addition, mtSSB decreases the fork regression kinetics (or pressure), which in turn, increases the probability of finding the holoenzyme at the moving state from ∼4 to ∼12% (SI). Even in the presence of mtSSB, the equilibrium is shifted towards the pause or inactive-state strongly (*K*(0), Table 1). Destabilization of *∼*2 base pairs (*d*) of the DNA junction by application of mechanical tension (*f*) is required to shift the equilibrium towards the moving state. For all figures error bars show standard errors. Values at 0pN were obtained from bulk measurements.

**Figure 4:**
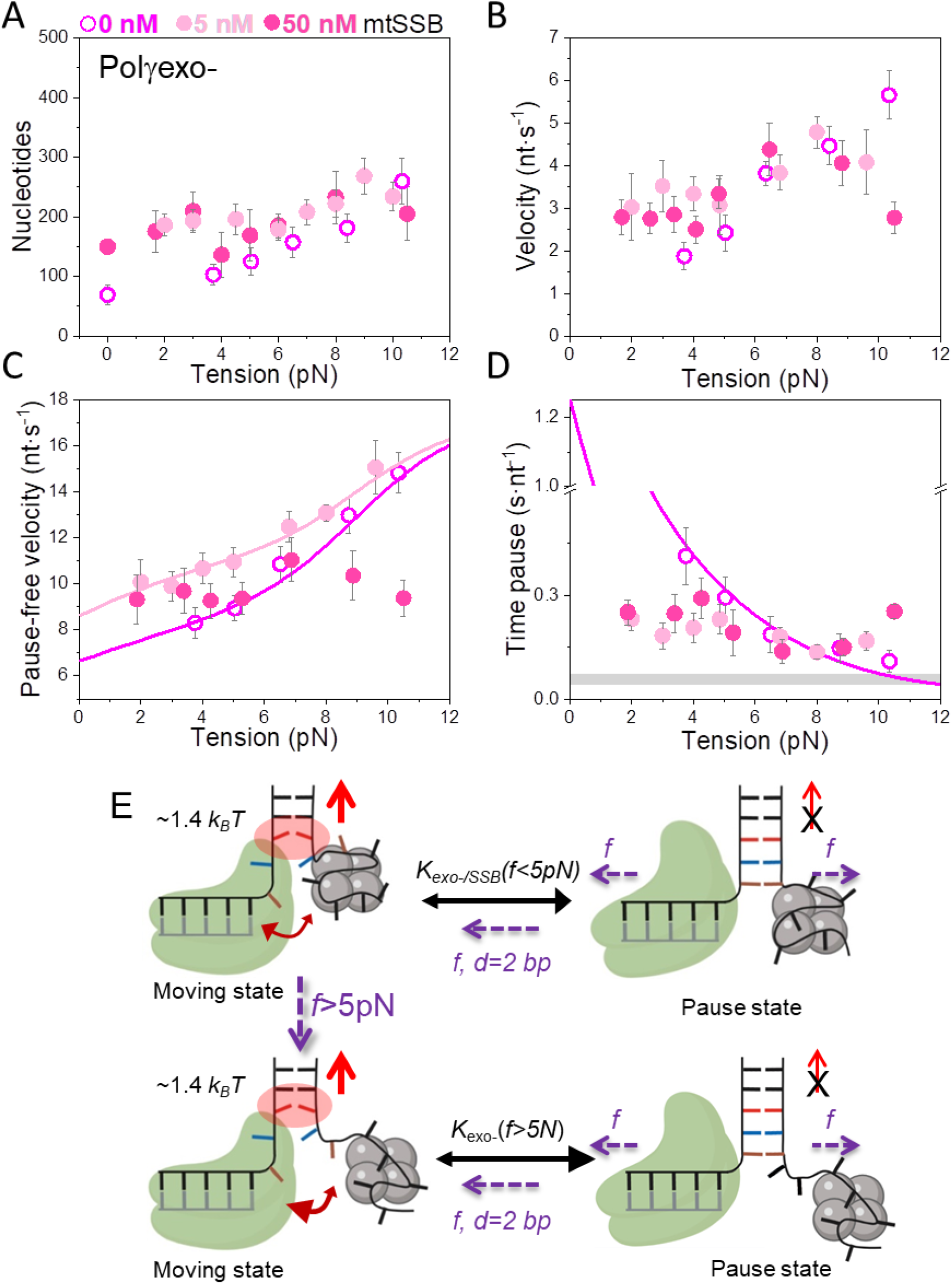
Effect of mtSSB on the tension dependent strand displacement kinetics of Polγexo-. Effects of 5 (N=44) and 50 (N=78) nM mtSSB on **A)** the average number of replicated nucleotides (0pN values correspond to bulk measurements), **B)** average strand displacement rate (nt s^-1^), **C)** Pause-free velocity, and **D)** average residence times at pause state per nucleotide (*T*_*p*_(*f*), s nt^-1^) of Polγexo-. Grey box shows average values obtained under primer extension conditions in the absence of SSB, Figure S1. Both mtSSB concentrations stimulate the strand displacement activity of Polγexo- below 4-6pN. However, the stimulatory effects diminish above 4-6 pN. Even more, as tension increased above ∼8 pN, 50 nM mtSSB had a detrimental effect on the average replication rates (**B**), pause-free rates (**C**), and residence time at pause state per nucleotide (**D**) of the mutant holoenzyme variant. In (**C**) magenta and pink lines correspond to the fits of the strand displacement model to the tension dependent pause-free rates in the absence and presence of 5 nM mtSSB, respectively. In (**D**) the magenta line is the fit of the two-state model (*Eq*.1) to the tension dependent average residence time at pause state per nucleotide in the absence of mtSSB. **E)** Diagram illustrating the effect of tension on the moving-pause state equilibrium of Polγexo- in the presence of SSB. At tension *f*<3-5pN, Polγexo- would alternate between moving and pause state with an equilibrium constant (*K*_*exo-/SSB*_(*f*<5pN)) ∼3-times smaller than that in the absence of SSB. Application of tension (f>3-5pN) to the DNA hairpin promotes the release of several ssDNA nucleotides from the SSB, which could decrease its ability to counteract the fork regression kinetics. Under these conditions, the mutant holoenzyme would alternate between moving and pause states with an equilibrium constant similar to that in the absence of SSB (*K*_*exo-*_(f>5pN)). In both situations, mtSSB (5 nM) binding energy and kinetics would help the holoenzyme to destabilize the first base pair of the DNA fork (Δ*G*_*int*_ ∼1.4 *k*_*B*_*T* per dNTP incorporated). Destabilization of *∼*2 base pairs (*d*) of the DNA junction by application of mechanical tension (*f*) will further shift the equilibrium towards the moving state in the two situations. For all plots error bars show standard errors.

We noted that the average and pause-free rates of Polγ and Polγexo- (with and without mtSSB) were identical in the presence and absence of T7DNAp in solution at all tensions, Figure S4. As reported for other DNA replication systems (24,57), these results indicate that polymerase exchange was not limiting and did not contribute significantly to the kinetics of the frequent pause events characteristic of the replication traces (Figures 1C, 1D and S3). This observation was further supported by additional results showing that, in the absence of T7DNAp, varying the concentrations of Polγ or Polγexo- by 20-fold did not alter their average rates and processivities significantly, Figure S5.

### Polγ and Polγexo- present identical fork destabilization energies

For both holoenzymes, the average processivities and velocities increased with tension, indicating that mechanical destabilization of the fork favors the strand displacement activity (Figure 2A and 2B). The strand displacement activity of Polγexo- was detected consistently at tension lower than that of Polγ (∼3 and 6 pN, respectively) and at all tensions presented faster average velocities than the wild-type holoenzyme, which is in agreement with the higher ability of the mutant variant to perform strand displacement synthesis in bulk (16,22,25,27). To further investigate the differences in the strand displacement activities of the two holoenzymes, we measured the effect of fork stability on the moving and pause states of their activities.

To examine the effect of fork stability on the moving state, we calculated the strand displacement rates without pauses, or pause-free velocity, at all tensions, Figure 2C (Methods). For both polymerases, pause-free velocity increased with tension similarly towards values found during primer extension (Figures 2C and S1C), indicating that DNA unwinding is the rate limiting step of the reaction. Pause-free velocity was ∼12% faster for Polγexo- than for Polγ, in agreement with the difference between the maximum primer extension rates of each holoenzyme (Figure S1C). Next, we fit the force dependent pause-free velocity of each holoenzyme to the theoretical framework described by Betterton and Julicher to quantify the unwinding activeness of nucleic acid helicases adapted to the case of replicative DNA polymerases (SI) (28,67,70). From now on, we will refer to this model as the strand displacement model. According to this model, two variables determine the maximum strand displacement rate of a DNA polymerase: the interaction energy of the polymerase with the fork, Δ*G*_*int*_, and the range on this interaction, *M*. These two free-parameters were fixed by least squares fits of the model to the pause-free velocity values of Polγ and Polγexo-, Figure 2C. The fits yielded Δ*G*_*int*_= 0.9±0.1 *k*_*B*_*T* and *M*= 1 for Polγ, and Δ*G*_*int*_= 1.0±0.1 *k*_*B*_*T* and *M*= 1 for Polγexo- (Table 1). These results indicated that both holoenzymes decrease the activation energy of the nearest bp of the fork equally by ∼1 *k*_*B*_*T*. Therefore, other factors should account for the different ability of each holoenzyme to perform strand displacement DNA synthesis.

### Polγ spends longer times in a non-productive state than Polγexo-

Next, we studied the effect of fork stability on the pause state of each holoenzyme by quantifying the effect of tension on their average residence times in pause state per nucleotide, *T*_*p*_(*f*) (Methods). Note that *T*_*p*_(*f*) includes pause frequency and duration. The results showed that *T*_*P*_*(f)* was higher for Polγ than for Polγexo- and for both holoenzymes decreased exponentially with tension towards values found during primer extension (Figure 2D). Previous studies on strand displacement DNA synthesis proposed that reannealing of the newly unwound bases would pause polymerase advancement by competing for template binding and promoting partition of the primer end from the *pol* to the *exo* site (24,28,71). In the case of Polγexo-, the frayed primer end would bind to the inactive *exo* site before returning intact to the *pol* site intramolecularly and favoring *pol* activity (11). Conversely, in Polγ, the newly incorporated nucleotide would be removed at the active *exo* site upon primer partition, rendering the holoenzyme prone to idle at the fork in recurrent *pol* and *exo* events (16). We note that we did not detect *exo* events of Polγ at any tension, suggesting that this reaction, and/or associated idling, involves few nucleotides not resolved by our current resolution limit. Therefore, although different in nature, the events triggered by the fork regression pressure on each holoenzyme would be detected as pauses under our experimental conditions. In a simplified two-state scenario, in which each holoenzyme alternates between a moving and a pause states (Figure 2E), the effect of tension on the average residence time in pause state per nucleotide during strand displacement can be quantified as (SI):

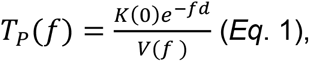

where, *K*(0) is the equilibrium constant of the transition between moving and pause state during strand displacement in the absence of tension. *f* is the mechanical tension that favors unwinding of the hairpin. *d* is the tension-induced conformational change along the pulling coordinate that shifts the equilibrium towards the moving state and *V*(*f*) is the tension dependent pause-free velocity defined by the strand displacement model described above (SI). The two free variables, K(0) and *d*, were fixed upon least-squares fitting of *Eq.1* to *T*_*p*_(*f*) data, Figure 2D and Table 1.

Extrapolation of the fits to 0 pN revealed that the average residence times at pause state per nucleotide in the absence of tension, *T*_*p*_(0), of Polγ and Polγexo- were 3.10 s nt^-1^ and 1.22 s nt^-1^, respectively (Table 1). These *T*_*p*_(0) values were ∼30 and ∼10 times higher than those during primer extension (∼0.1 s nt^-1^ for both holoenzymes (17)), Figures 2D and S1D, showing that stability of the DNA fork has a strong effect on increasing Tp(0). In fact, this effect is higher than that on decreasing pause-free rates of each holoenzyme. In terms of moving probabilities, the *T*_*p*_(0) values indicate that the probabilities of finding Polγ and Polγexo- moving through the DNA hairpin are as low as ∼4 and ∼12%, respectively (Methods, SI). This data is in line with the stronger ability of the mutant variant to perform strand displacement DNA synthesis (16).

The value of the tension-induced conformational change that shifts the equilibrium towards the moving state obtained from the fits was *d* ∼1.2 nm for the two holoenzymes. This distance is compatible with the gain of ∼4 single-stranded nucleotides along the pulling coordinate (average extension per nucleotide at *f*<11 pN, ∼0.26 nm nt^-1^, (17)). This result is in agreement with a mechanism in which destabilization of the ∼2 first bp of the DNA fork by tension diminishes the fork regression pressure and creates two template nucleotides that could be accommodated at the polymerase template-binding pocket (72) shifting equilibrium towards the moving state, Figure 2E.

Finally, we note that the values of *V*(0).and *T*_*p*_(0) predicted by fits to the single molecule data (Figures 2C and 2D), together with the residence times at the DNA fork of each holoenzyme determined in bulk (Table 1), imply average processivities in the absence of tension of ∼57 nt and ∼92 nt, for Polγ and Polγexo- respectively (Methods). Given the differences between bulk and single-molecule approaches, these values could be considered in line with the maximum number of replicated nucleotides measured in bulk studies, 37±15 for Polγ and 69±17 for Polγexo- (Figure 1E, Table 1), supporting the models used to explain the data.

### Differential effects of mtSSB on Polγ and Polγexo- strand displacement activities under mechanical tension

Next, we aimed to investigate the effect of mtSSB interaction with the displaced strand on the tension dependent strand displacement replication kinetics of Polγ and Polγexo- variants. We tested the effects of 5, 50 and 100 nM mtSSB, which under our current experimental conditions have been shown to cover respectively ∼85%, ∼98% or are expected to oversaturate individual ssDNA molecules stretched under mechanical tension (51). In the case of Polγ, 5 nM mtSSB favored detection of activities at slightly lower tensions (∼4-5 pN), increased the average rate without pauses, and decreased *T*_*p*_(f) by 2-3-fold with respect to conditions in the absence of mtSSB, but did not alter pause-free rates, Figures 3A-D. In contrast, 5 nM mtSSB had stronger stimulatory effect on the strand displacement activity of Polγexo-; i.e., it dropped the minimum tension required to detect individual activities, from ∼3 to ∼1 pN, increased the average processivity, average replication rate and the pause-free velocity (by ∼25%), and decreased the time at the pause state by ∼4 times, with respect to the values predicted by the strand displacement and the two-state models for conditions in the absence of SSB (Figures 4A-D). Interestingly, while the stimulation of pause-free velocity of the mutant holoenzyme by 5 nM mtSSB persisted at all tensions, the stimulation of the average rate, processivity and residence time at pause state decreased with tension above ∼2-3 pN to ceased at ∼5 pN (Figures 4A-E).

Increasing mtSSB concentration to 50 nM promoted the strand displacement activity of Polγ strongly; it dropped the minimum tension required to detect individual activities of from ∼6 to ∼3 pN, stimulated the average processivity, the average rates and pause-free rates (∼25% increase), and decreased the time at pause state per nucleotide (∼2-3 times) at all tensions with respect to conditions in the absence of mtSSB (Figures 3A-D). In the case of Polγexo-, the stimulatory effects of 50 nM mtSSB on the strand displacement activity were identical to those measured for 5 nM mtSSB and again, only observed at the lowest tension (<5 pN), Figure 4. Notably, under these conditions, as tension increased above 6-8 pN, the average replication rates (with and without pauses) were significantly slower and time at pause state higher than those in the absence of mtSSB, showing a detrimental effect of this mtSSB concentration on Polγexo- activity at high tensions (Figures 4B-D). Further increase of mtSSB concentration to 100 nM resulted in lack of stimulation and further inhibition of Polγ and Polγexo- strand displacement activities Figures S6 and S7. These results are in line with previous bulk biochemical assays that showed deleterious effects of oversaturating mtSSB concentrations on the DNA synthesis activity of the human mitochondrial holoenzyme (42). Lastly, we note that, at the lowest detection tensions, mtSSB (5 and 50 nM) did not alter significantly the exchange rates, *k*_*exc*_ (calculated as the ratio between average velocity and processivity), of Polγ and Polγexo- with competing T7DNAp in solution (Table 1).

Overall, the above results showed that mtSSB stimulates the strand displacement of both holoenzymes by favoring detection of strand displacement activity at lower forces, increasing pause-free rates and decreasing the time at non-productive or pause state. Interestingly, the real-time kinetics of each holoenzyme responded to the combined effect of mtSSB concentration and tension differently, Polγexo- being more prone to stimulation by lower mtSSB concentrations (10-fold) but also to inhibition by tension in the presence of higher mtSSB concentrations.

### Effects of noncognate EcoSSB and gp2.5 proteins on Polγ and Polγexo- strand displacement activities under mechanical tension

To determine whether the observed effects of mtSSB on the strand displacement replication by Polγ depend on specific interactions between these two factors, we assessed the effects of tension and concentration of homologous EcoSSB and heterologous phage T7 SSB, gp2.5, on the average rates (with and without pauses) and times at pause state of each Polγ variant. These experiments were performed in the absence of competitor T7DNAp in solution because EcoSSB and gp2.5 stimulated the strand displacement activity of the phage polymerase (see Figure 6), which in turn, hindered the detection of polymerase exchange at the lowest tensions. We note again that polymerase exchange did not affect average rates and time at pause state of Polγ and Polγexo- (Figure S4), validating the comparison of these data with those taken in the presence of competitor T7DNAp.

Bulk biochemical assays show that saturating concentrations of EcoSSB stimulated the strand displacement processivity of Polγ and Polγexo- in a way similar to that measured for its mtSSB homolog (Figure S2, Table 1). However, notable differences were apparent at the single-molecule level between the effects of the two SSB proteins on the real-time kinetics of each holoenzyme. At the lowest concentration (5 nM), EcoSSB did not have significant effects on the initiation force, tension dependent average rates (with and without pauses) and times at pause states of both holoenzymes (Figures 5A-F). These results contrast with the stimulatory effects of 5 nM mtSSB on the strand displacement kinetics of the two holoenzyme variants (Figures 3 and 4). The increase of EcoSSB concentration to 50 nM stimulated the activity of the two holoenzymes similarly (Figures 5A-F); i.e., the minimum tensions required to detect activities decreased (to ∼3 and ∼1 pN for Polγ and Polγexo-, respectively), and the average rates of the two holoenzymes at the lowest tensions were stimulated by increasing the pause-free velocities (∼25% increase) and decreasing the time at pause state per nucleotide (2-4 times). In sharp contrast with mtSSB (50 nM), EcoSSB (50 nM) did not present inhibitory effects on Polγexo- activity at high tensions and stimulated the pause-free velocities of both holoenzymes at all tensions (Figures 5B and 5E). However, EcoSSB (50 nM) lost the ability of increasing the average rate and/or decreasing *T*_*p*_(*f*) of Polγ at tension above 5-6 pN (Figure 5C). As in the case of high mtSSB concentrations, no stimulation or inhibition of the strand displacement activities of both holoenzymes was measured with 100 nM EcoSSB at high tensions (*f*> 8pN), Figure S8.

**Figure 5:**
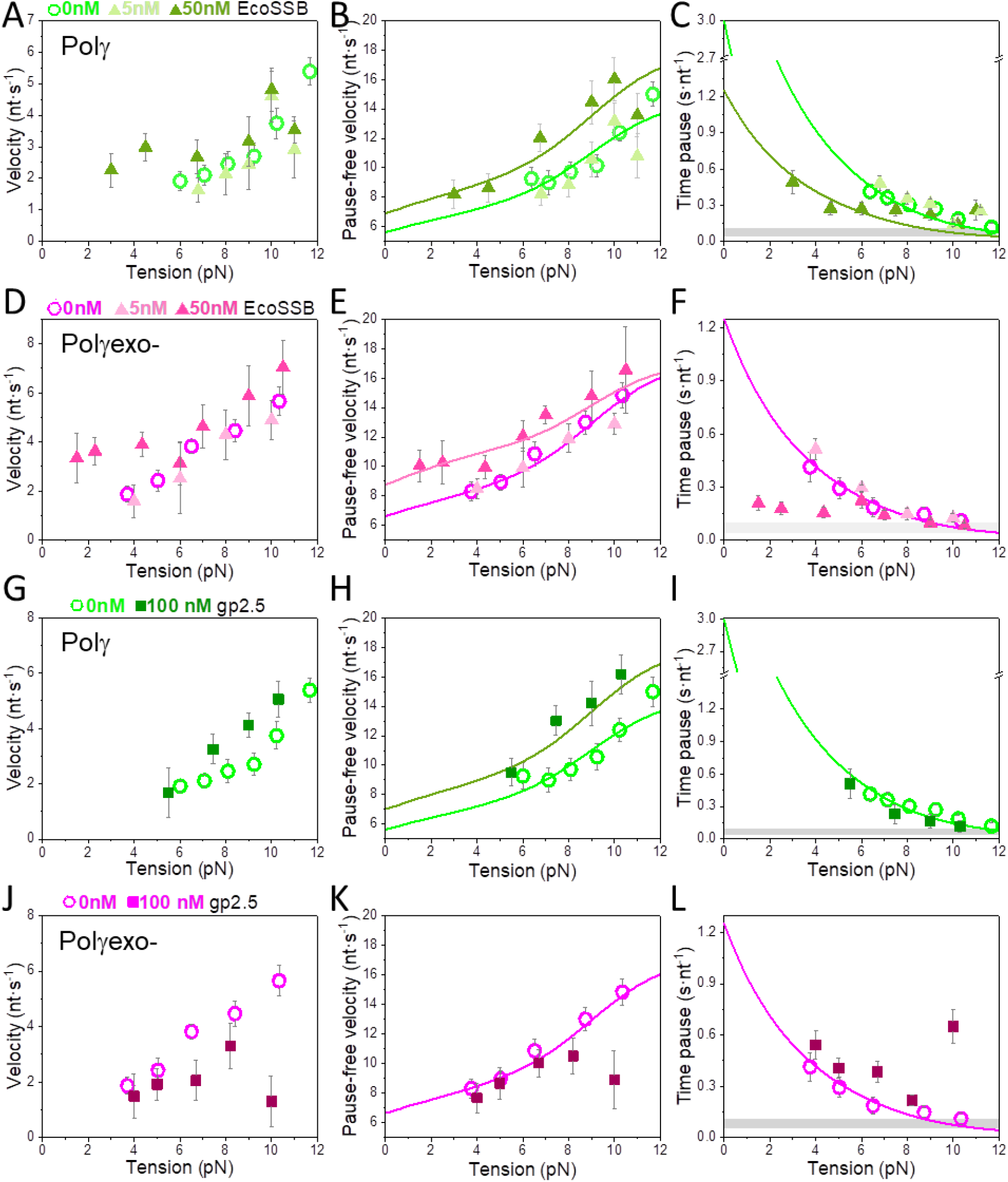
Effects non-cognate SSB on Polγ and Polγexo-tension dependent strand displacement kinetics. **A), B)**, and **C)** 5 nM EcoSSB had no apparent effect on the strand displacement kinetics of Polγ (N=28). In contrast, 50 nM EcoSSB (N=32) stimulated the average rates (**A**), pause-free velocity (**B**), and residence time at pause state (**C**) of Polγ below 5 pN. Whereas the stimulation of pause-free velocity was force independent, the stimulatory effects of 50 nM EcoSSB on the average rate and residence time at pause state disappear at tension above 5 pN. **D), E)**, and **F)** 5 nM EcoSSB (N=27) had no significant effects on the strand displacement kinetics of Polγexo-, while 50 nM EcoSSB (N=42) stimulated the average rates (**D**), and time at the pause state (**F**) below 5 pN, and the pause-free velocity (**E**) at all tensions. **G), H)**, and **I)** gp2.5 (100 nM, N=15) stimulated the pause-free velocities of Polγ to a similar extend than 50 mM mtSSB and EcoSSB (**H**), but did not decrease the residence time at pause state of the wild-type variant at any tension (**I**). **J**), **K**), and **L**) gp2.5 (100 nM, N=17) did not stimulate the strand displacement kinetics of Polγexo- and was inhibitory at tension above ∼8 pN. Light and dark green lines correspond to the fits of the strand displacement model (**B, H**) and *Eq*. 1 (**C, I**) to Polγ data in the absence and presence of mtSSB (50 nM), respectively. Magenta and pink lines are the fits of the strand displacement model (**E, K**) and *Eq*. 1 (**F, L**) to Polγexo- data in the absence and presence of mtSSB (5 nM), respectively. Grey boxes show the average *T*_*p*_(*f*) values obtained during primer extension conditions in the absence of SSBs (Figure S1 and (16)). For all plots error bars show standard errors.

In the case of non-cognate heterologous gp2.5, bulk biochemical assays showed that the phage SSB stimulated the processivity of Polγ but not that of Polγexo- (Figure S2A). At the single-molecule level, only the highest gp2.5 concentration used in our experiments, 100 nM, had significant effects on the activity of the two holoenzymes, probably reflecting the lower affinity of this protein for ssDNA. Under these conditions, gp2.5 did not favor the detection of Polγ and Polγexo- activities at tensions significantly lower than those in the absence of SSB (Figures 5G to 5L). However, gp2.5 (100 nM) stimulated the pause-free velocities of Polγ to the extent similar as in the cases of 50 mM mtSSB and EcoSSB, but did not decrease the residence time at pause state of the wild-type variant at any tension, Figures 5G-I. In contrast, gp2.5 (100 nM) did not stimulate and even inhibited the strand displacement activity of Polγexo- (especially at high tensions), Figures 5J-L. These results showed again that the strand displacement activity of Polγexo- is more sensitive to the combined effect of tension and SSB than Polγ.

Although effects of SSBs on the two holoenzymes at hand differ in details, the results generally showed that at concentrations ∼10 to 20 times lower than those of noncognate SSBs, mtSSB has greater ability to stimulate the strand displacement kinetics of the two Polγ variants under mechanical tension. Notably, the lack of stimulation and/or inhibition of Polγexo- average strand displacement rate by the three SSBs under study at tensions above ∼5 pN, suggest that under mechanical stress conditions the mutant holoenzyme cannot correctly couple with these SSB proteins during strand displacement replication.

### Effects of SSBs on the strand displacement activities of T7 DNA polymerase variants under mechanical tension

Finally, we sought to determine whether our observations of the behavior of mitochondrial holoenzymes in response to various SSB proteins and tension can be extrapolated to other polymerase-SSB systems. To this end, we measured the effects of cognate (gp2.5) and non-cognate (mtSSB and EcoSSB) SSBs on the kinetics of the strand displacement replication by the wild-type (T7DNAp) and the *exo*-deficient (Sequenase©) variants of phage T7 DNA polymerase under increasing mechanical tensions. T7DNAp shares sequence, structural and functional homology with the catalytic subunit of Polγ, PolγA (73).

The potent *exo* activity of T7DNAp (Figure 6A), prevented the detection of strand displacement activities below ∼8 pN, as reported previously (24). In sharp contrast, the absence of *exo* activity in Sequenase© (Figure 6A) allowed the detection of strand displacement actives of this variant at tensions ∼4 pN. Upon exclusion of *exo* events from T7DNAp traces, the two phage polymerase variants presented almost identical tension-dependent average rates (with and without pauses), and residence times at the pause state per nucleotide (Figures 6C-H). This similarity is likely due to the effective identification and removal of *exo* events, which do not longer contribute to the pause state kinetics, as in the case of Polγ data. In fact, least-square fits of Sequenase© pause-free and *T*_*p*_(*f*) data to the strand displacement and two-states model (*Eq*.1), respectively, explained well the corresponding values of T7DNAp (Figures 6D, 6E, 6G and 6H). On the one hand, fits of pause-free velocity data yielded Δ*G*_*int*_= 0.3±0.1 *k*_*B*_*T* (*M*= 1) per incorporated nucleotide (Figures 6D and 6G, Table S2). These results indicate that the phage polymerase destabilizes the first bp of the DNA fork with energy ∼2-3 times weaker than that measured for the mitochondrial counterpart. On the other hand, fits of residence times at pause state data (*T*_*p*_(*f*)) with *Eq*.1 yielded the values of *K(0)*=3.52 and *d*_*T7*_ ∼0.84 nm and predicted that in the absence of tension *T*_*p*_(*0pN*)= 0.32 nt s^-1^ (Figures 6E and 6H, Table S2). As found in the case of Polγ, the value of *T*_*p*_(*0pN*) for the phage DNApol is ∼30 times higher than that during primer extension conditions (∼0.01 nt s^-1^, Figure 6E and (17)), and again, the strong negative effect of fork stability on pause kinetics is overcome by the mechanical destabilization of the ∼2 first bp of the fork (*d*_*T7*_∼0.84 nm).

**Figure 6:**
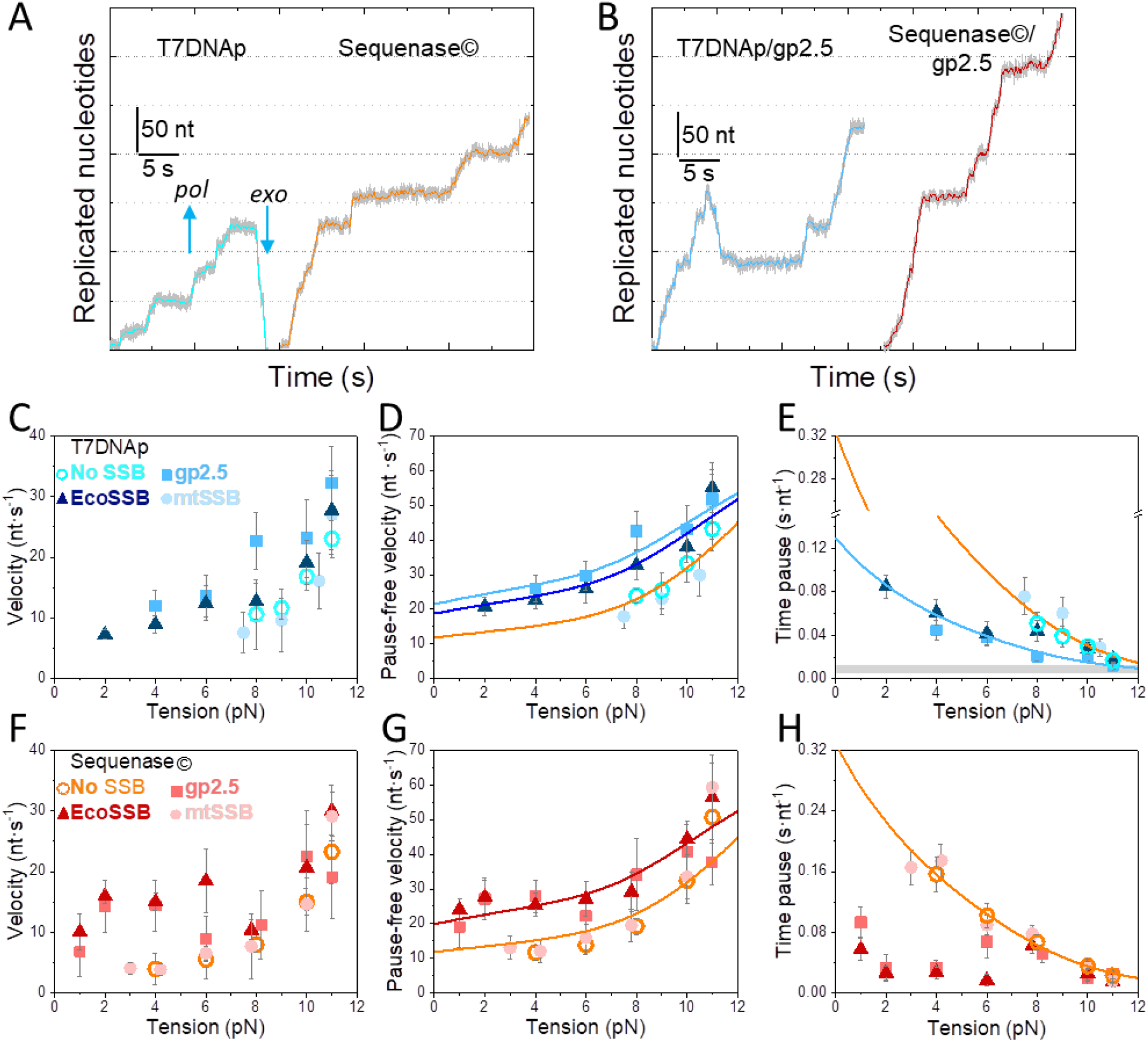
Effect of cognate and non-cognate SSBs on T7DNAp and Sequenase© tension dependent strand displacement kinetics. Representative strand displacement traces of T7DNAp and Sequenase© in the absence **A**) and presence **B**) of gp2.5 (*f*∼8 pN). T7DNAp showed processive *exo* events in the absence and presence of SSBs. **C**), **D**) and **E**) tension effects on the T7DNAp (N=34) average rate (**C**), pause-free rate (excluding *exo*) (**D**), and residence time at the pause state per nucleotide (**E**). gp2.5 (100 nM, N=31) increased average rate (**C**) and pause-free rate (**D**), and decreased times at the pause states (**E**) of T7DNAp at all tensions with respect to the values in the absence of SSB. EcoSSB (50 nM, N=23) stimulated pause-free velocity at all tensions (**D**), and the average rates (**C**) and residence time at pause state (**E**) at tension below 8 pN. **F), G)** and **H)** effects of tension on Sequenase© (N=27) average rate (**F**), pause-free rate (**G**) residence time at the pause state per nucleotide (**H**). gp2.5 (100 nM, N=23) and EcoSSB (50 nM, N=24) stimulated the pause-free velocity at all tensions (**G**) and the average rates (**F**) and residence time at pause state (**H**) at tensions below 6-8 pN. In contrast, mtSSB (50 nM) did not have significant stimulatory effects on the strand displacement activities of T7DNAp (N=17) and Sequenase© (N=31) (**C** to **H**). In (**D**) light and dark blue lines are the fits of the strand displacement model to T7DNAp tension dependent pause-free rate data in the presence of gp2.5 and EcoSSB, respectively. Orange line shows the fits with the strand displacement model to the pause-free rate of Sequenase© in the absence of SSB. In (**E**), light blue and orange lines are the fits of *Eq*. 1 to T7DNAp and Sequenase© data, respectively, and grey box shows position of the average *T*_*p*_(*f*) values measured under primer extension conditions (16). In **G**) orange and red lines correspond to the fits with the strand displacement model to the pause-free rate of Sequenase© data in the absence and presence of gp2.5, respectively. In **H**) orange line is the fit of Eq. 1 to the tension dependent residence time in pause state of Sequenase© in the absence of SSB. For all plots error bars show standard errors.

As measured for the mitochondrial polymerase, SSB stimulation of strand displacement kinetics of the phage DNA polymerases depended on tension and showed significant differences between each polymerase-SSB pair. In the case of T7DNAp, cognate gp2.5 (100 nM) *i*) favored the detection of strand displacement activities at tensions lower than those in the absence of SSB (∼3-4 pN), *ii*) increased the average replication rates and pause-free velocity, and *iii*) decreased the residence time at the pause state per nucleotide at all tensions, with respect to the values predicted by the strand displacement and the two-state models in the absence of SSB (Figures 6C-E). In the case of Sequenase©, gp2.5 (100 nM) also favored detection of activities at lower tension (∼1 pN) and stimulated pause-free velocity at all tensions, but failed to stimulate the average rate (or to decrease time at pause state) at tensions above 6pN (Figures 6F-H). These tension-dependent stimulatory effects resemble those of cognate mtSSB on the strand displacement activities of Polγ and Polγexo-. Addition of non-cognate EcoSSB had similar effects on the activities of T7DNAp and Sequenase© as a function of tension; EcoSSB (50 nM) stimulated pause-free velocity at all tensions, whereas it only stimulated the average rate (or decreased time at pause state) at tensions below 8 pN (Figures 6C-H). Interestingly, these effects are identical to those measured for the mitochondrial holoenzymes at this EcoSSB concentration. Finally, mtSSB (50 nM) did not have significant stimulatory effects on the strand displacement activities of T7DNAp and Sequenase©, Figures 6C-H. These results contrast with the multiple stimulatory effects of mtSSB on their cognate holoenzyme variants.

Overall, the effects of cognate and non-cognate SSBs on the tension dependent kinetics of strand displacement replication by T7DNAp and Sequenase© are in line with those measured on the mitochondrial holoenzymes. In summary, the results show that *i*) binding of SSBs to ssDNA alone is not sufficient to promote pause-free velocities, *ii*) cognate but not non-cognate SSBs decrease the residence time in non-active state of wild-type variants at all tensions, while *iii*) both cognate and non-cognate failed to decrease the residence times at the pause state characteristic of the *exo* deficient variants at high tensions. The implications of these results are discussed further.

### Quantification of SSB effects on strand displacement kinetics

Under conditions favoring strand displacement replication, cognate and non-cognate SSBs promoted the pause-free velocities of the mitochondrial and phage polymerase variants to a similar extent, by ∼20-25%. We fitted these data sets to the strand displacement model to quantify the energetic contribution of SSBs to DNA unwinding. In the case of the mitochondrial holoenzymes, least square fits yielded similar values of the DNA fork interaction/destabilization energies of Δ*G*_*int*_∼ 1.4 *k*_*B*_*T* (*M*= 1) for all holoenzyme-SSB couples (Table 1): Polγ/50nM mtSSB (Figure 3C), Polγexo- /5nM mtSSB (Figure 4C), Polγ/50nM EcoSSB (Figure 5B), Polγexo- /50nM EcoSSB (Figure 5E), and Polγ/gp2.5 (Figure 5H). Whereas for the phage variants, least square fits yielded Δ*G*_*int*_ ∼0.7 *k*_*B*_*T* (*M*= 1) in all cases (Figure 6D and 6G, Table S2). These results imply that, under conditions that allow stimulation, cognate and noncognate SSBs contribute non-specifically an extra ∼0.4 *k*_*B*_*T* to decrease the activation barrier of the first bp of the fork during the strand displacement activities of both the mitochondrial and phage polymerases.

To quantify the contributions of SSBs to decrease the residence times at the pause state(s) (T_p_(*f*)), we fitted to the two-state model (*Eq*.1) the only two cases in which the SSBs decreased *T*_*p*_(*f*) at all tensions: Polγ and T7DNAp in the presence of their cognate SSBs (Figures 3D and 6E). The fits yielded the equilibrium constant between pausing and polymerization (K(0)) and the magnitude of the conformational change that shifts equilibrium towards moving state (*d*) (Tables 1 and S2). For the two polymerases, extrapolation of the fits to 0 pN indicated that the presence of their cognate SSB decreased their respective average residences time at pause state in the absence of tension by ∼2-2.5 fold. On the other hand, the values of *d* that resulted from the fits were similar in both cases, *∼*1-1.2 nm (Tables 1 and S2). This conformational change is similar to that measured in the absence of SSB suggesting that in the presence of SSB, unwinding of the first ∼2 bp of the DNA fork by tension is still necessary to shift the equilibrium towards moving state. Note that these fits explained well the effects of non-cognate EcoSSB (50 nM) on the *T*_*p*_(*f*) of Polγ and T7DNAp at tensions below ∼6 and 8 pN, respectively, Figure 5C and 6E. Above these tensions, EcoSSB had no longer an effect on the *T*_*p*_(*f*) of both DNA polymerases. Similarly, the ability of cognate and non-cognate SSB to decrease the residence time at the pause states of *exo* deficient variants (Polγexo- and Sequenase©), was eliminated by increasing tension (Figures 4D, 5F and 6H). This effect limited the data points available for consistent fits with the two-state model. Nevertheless, the data at the lowest tension (∼1 pN) showed that cognate and non-cognate SSBs lowered the *T*_*p*_(1*pN*) of the two *exo* deficient variants ∼4-fold, with respect to the values predicted by the fit of the two-state model to data in the absence of SSB. This stimulatory effect is about two-fold higher than that measured for the same concentrations of SSB on the *T*_*p*_(0) of the wild-type variants.

## DISCUSSION

Replicative DNA polymerases often couple DNA synthesis with strand displacement or dsDNA unwinding activity. This capability facilitates the synthesis of the leading and the lagging DNA strands by displacing one of the parental DNA strands enabling the recruitment of SSB proteins onto the replication fork. SSB proteins assist and coordinate the activities of the polymerase with those of other protein partners at the DNA fork (13,40,74). Hence, studying strand displacement activity in the context of SSBs is essential to define this reaction more precisely and how these two interacting partners couple their activities at the DNA fork. Our results provide further insights into the mechanism of SSB-assisted strand displacement DNA synthesis of the human mitochondrial DNA polymerase.

Our data demonstrates that the human mitochondrial DNA polymerase, Polγ, and its *exo* deficient variant D_198_AE_200_A, Polγexo-, present consistent strand displacement DNA synthesis in bulk and single-molecule optical tweezers assays. Single-molecule measurements showed that during this reaction both holoenzymes exchanged at the fork with competitor T7DNAp polymerases in solution. The polymerase exchange rate was not limiting and did not contribute to the pause kinetics. Similarly, single-molecule and bulk studies on prokaryotic and eukaryotic replisomes have shown fast (non-limiting) polymerase exchange within active DNA replication both *in vivo* and *in vitro* (56,57). Fast exchange of the mitochondrial holoenzymes at the DNA fork likely occurs also in the absence of competitor T7DNAp in our optical tweezers experiments, explaining why the average processivity of the replication traces at the lowest tension were higher (2-3 times) than those expected from the single turn over bulk experiments (Figure 1E). In addition, we did not detect consistent effects of mtSSB on the apparent dissociation and exchange rates of either holoenzyme from the DNA (Table 1). This argues that the stimulatory properties of mtSSB result from effects distinct from those of increasing the residence time of the polymerase on DNA (see below).

Upon initiation, the two mitochondrial holoenzymes decreased the activation energy of the nearest bp of the fork equally, by Δ*G*_*int*_ ∼1 *k*_*B*_*T* per dNTP incorporated, suggesting that the intrinsic strand displacement mechanism of Polγ is not interfered by its *exo* activity. According to current models of the mechanism of coupling DNA synthesis with unwinding, the sharp bending of template (∼90°) induced characteristically by DNA polymerases within their polymerization domains (72,75) would impose mechanical stress at the DNA fork junction, which in turn lowers the energy barrier for DNA unwinding during each nucleotide incorporation cycle, Figure 2E (24,28,32,67). Under our experimental conditions, the Δ*G*_*int*_ of the mitochondrial holoenzyme is ∼45% lower than the average stability of the next bp of the hairpin, Δ*G*_*bp*_(0pN) ∼1.8 *k*_*B*_*T* (SI). This difference explains why, in the absence of tension, the activation energy of fork melting decreases ∼4 times the pause-free velocity of both holoenzymes with respect to that during primer extension conditions in the absence of secondary structure, ∼6 vs ∼24 nt s^-1^, respectively. Regardless, under these conditions, the pause-free nucleotide incorporation rate would still be ∼120 times faster than the rate of partitioning of a correctly base-paired primer from the *pol* to the *exo* domain (11). Therefore, as long as the template is bound to *pol* site, despite lowering the replication rate, the fork stability will not promote the kinetic partitioning of the primer from the *pol* to the *exo* sites, which otherwise would interfere with forward movement of the holoenzyme.

What is the main factor limiting the strand displacement activity? Previous single-molecule studies suggested that the fork regression pressure outcompetes the *pol* active site for binding to the template and consequently impedes the forward translocation of the enzyme, Figure 2E (24,28,71). These conditions would favor the kinetic partitioning of the primer from the *pol* to *exo* sites, We propose that this process underlays the extended residence time each holoenzyme spends in pause or non-productive states (*T*_*p*_(*f*)), which resulted in probabilities as low as ∼4 and ∼12% of finding Polγ and Polγexo- moving through the DNA hairpin in the absence of tension, respectively (SI). According to our data *T*_*p*_(0) are 30 and 10 times higher (for Polγ and Polγexo-, respectively) than those during primer extension conditions and therefore, the main factor limiting the elongation phase of the strand displacement replication In the case of the *exo* deficient mutant, the frayed primer end would shift to the inactive *exo* site before returning intact to the *pol* site (10,11). Conversely, the wild-type Polγ would idle at the DNA fork junction in a process in which a new incorporated nucleotide is removed at the active *exo* site upon the intramolecular transfer of the primer between *pol* and *exo* sites (16). Cyclic idling cannot be resolved with our current resolution limit and would be added to the pause kinetics of wild-type Polγ, explaining why it spent 2-3 times longer residence times at the pause state per nucleotide than the *exo* deficient variant. In agreement with this hypothesis, effective identification and removal of the fast *exo* activities from replication traces of T7DNAp showed that the wild-type and *exo* deficient phage polymerases spent similar average residence times at pause state per nucleotide. Subsequently, the average velocity of Polγ is 2-3 times slower than that of Polγexo-, which explains the higher processivity (2-3 fold) of the former on dsDNA (note that the two enzymes have similar dissociation rates from DNA). According to our 2-state model (*Eq*.1), destabilization of the first ∼2 bp of the fork by application of mechanical tension to the DNA hairpin would release fork regression pressure on the mitochondrial (and phage) holoenzymes and create 2 ssDNA nucleotides for their template-binding pockets shifting the equilibrium towards productive DNA synthesis, Figure 2E. These results are in agreement with previous studies showing two ssDNA template nucleotides bound at the polymerization domain of the human mitochondrial and phage T7 replicative DNA polymerases (72,75,76).

According to our data, cognate and non-cognate SSBs stimulate the processivity of the strand displacement activities of Polγ and Polγexo- by increasing the pause-free rates and decreasing the residence time at pause state per nucleotide (i.e., increasing average replication rates). However, these stimulatory effects were evident only under specific conditions. On the one hand, cognate and non-cognate SSBs stimulated pause–free velocity a 25% by decreasing the energy barrier for the unwinding of the first bp of the DNA fork, by additional ∼0.4 *k*_*B*_*T* (Figure 3E). This contribution is in excellent agreement with the average binding energies per nucleotide measured for mtSSB and EcoSSB under similar experimental conditions (51,77). Overall, stimulation of pause-free velocities depended on SSB concentration but not on the external mechanical tension exerted to the complementary strands of the hairpin. Previous single-molecule manipulation studies showed that mechanical tension applied to tetrameric SSB-ssDNA complexes affects the SSB binding footprint on ssDNA (77). Taken together, these results suggest that the binding energy and kinetics but not the binding footprint (wrapping mode) of the SSB to/on ssDNA could be relevant to stimulate pause-free velocities. In addition, we note that 5 nM mtSSB but not 5 nM EcoSSB or 50 nM gp2.5 stimulated the pause free velocity of Polγexo- at all tensions, and gp2.5 (100 nM) and EcoSSB (50 nM) but not mtSSB (50 nM) promoted the pause-free rates of the phage T7DNAp and Sequenase© variants. These results indicate that species-specific interactions between the polymerase and the SSB may also play a role in the coordination of the polymerase and SSB activities at the fork and promotion of pause-free rate of strand displacement DNA synthesis. It is tempting to speculate that these interactions would be likely functional in the case of the mitochondrial polymerase-mtSSB system (41) and physical between the phage holoenzyme and the C-terminal tail of gp2.5 and EcoSSB (74,78). Noteworthy is that the specific interactions also seem to facilitate the inhibition of the pause-free rates of Polγexo- at high mtSSB concentration and tensions, showing the increased sensitivity of this variant to the cognate SSB.

On the other hand, under conditions that allowed stimulation at the lowest tension (see below), SSB interaction with the displaced strand decreased ∼2-4 times the residence times in the non-productive or pause state (*T*_*p*_(*f*)) of Polγ and Polγexo-, respectively. Under our experimental conditions none of the three SSBs under study presented in isolation DNA unwinding activities. However, additional single-molecule experiments showed that the three SSBs do have the ability to decrease the reannealing rate of the complementary strand of the hairpin dramatically (∼100-1000-fold, Figure S9). Therefore, it is tempting to speculate that the ability of SSB to reduce the fork regression kinetics (regression pressure), may decrease the competition between the DNA fork and the polymerase for binding to the ssDNA template (Figure 3E), thus decreasing the average residence time in pause state of the polymerase. Remarkably, the stimulatory effects of SSBs on *T*_*p*_(*f*) presented, in many cases, a strong dependency on tension. For example, the ability of cognate and noncognate SSBs to decrease *T*_*p*_(*f*) of Polγexo- and Sequenase© faded away as tension raised to ∼5 pN. As mentioned above, mechanical tension applied to SSB-DNA complexes diminishes the binding footprint of the SSB to ssDNA. This process occurs by the gradual unwrapping of ssDNA nucleotides from tetrameric SSBs as tension increases (51,77). Therefore, it is tempting to speculate that mechanical unwrapping of ssDNA from the SSB would lead to physical separation of the SSB from the fork junction, which in turn, would decrease its ability to counter act the fork regression pressure and control the moving-pause equilibrium of *exo*-deficient variants (Figure 4E). Interestingly, mtSSB and gp2.5 retained the ability to decrease *T*_*p*_(*f*) of their cognate wild-type polymerases (Polγ and T7DNAp, respectively) at all tensions, whereas noncognate SSBs did not. These results argue that specific polymerase-SSB interactions would help to control the wild-type Polγ pauses (which may include idling) even under mechanical stress conditions. In addition, our results showed that mtSSB decreased the *T*_*p*_(*f*) of the two mitochondrial holoenzymes at a concentration ten-fold lower than that of homologous EcoSSB, whereas noncognate gp2.5 had no significant effects on pause kinetics. These results support again that, even though no physical interactions have been described between Polγ-mtSSB (41), specific functional polymerase-SSB interactions could play a role in the modulation of the equilibrium between the moving and pause states characteristic of each polymerase.

We note that the effects of SSBs on *T*_*p*_(∼0 pN) were ∼2 times stronger than those on the pause-free velocities, pointing it out as the main factor that promotes the average velocity (and, thus, the average processivity) of the strand displacement activity, and favors the detection of activities of the mitochondrial holoenzyme variants at significantly lower tensions (≤3 pN). In any case, even in the presence of cognate mtSSB, the *T*_*p*_(∼0pN) of Polγ and Polγexo- were 10 and 4 times higher, respectively, than those during primer extension conditions, implying that their moving probabilities through the DNA fork in the absence of tension were restricted to ∼12 (for Polγ) and 30% (for Polγexo-) (SI). In addition, mechanical destabilization of the first ∼2 bp of the DNA fork was still required to shift the equilibrium away from pause state. These results argue that mtSSB has only moderate impact on shifting the equilibrium towards the active polymerization competent state of the holoenzyme.

Overall, our results show that human mitochondrial DNA polymerase holoenzyme exhibits a robust strand displacement DNA mechanism, which is further enhanced by the mtSSB. In the presence of mtSSB, Polγ and Polγexo- variants synthetize hundreds of nucleotides through the fork. In the case of Polγexo-, even sub-saturating mtSSB concentrations (5 nM) were sufficient to boost the intrinsic strand displacement activity of the mutant holoenzyme variant. The functional interactions between Polγ and mtSSB likely have significant physiological implications. On the one hand, because strand displacement activity is inversely related to polymerase slippage (13), the potent strand displacement activity of Polγ would generally favor its fidelity by decreasing the probability of deletions (and/or insertions) during replication of the lagging or L-strand of the mtDNA. On the other hand, reports to date indicate that ∼95% of all the mtDNA synthesis events initiated at the origin of replication (i.e., O_H_) are terminated prematurely at the termination associated sequence (TAS), likely due to the absence of mtDNA helicase Twinkle (29). This observation implies that for the vast majority of time, Polγ resides at the fork junction (the end of the D-loop, near TAS) accompanied only by the displaced strand-bound mtSSB, which resembles our study model. It is tempting to speculate that the strand displacement activity is exerted at TAS and perhaps serves the purpose of providing an optimal substrate for Twinkle to bind and complete the replisome, initiating this way the processive replication of mtDNA. The occurrence of strand displacement mtDNA synthesis at TAS under physiological conditions is advocated by the fact that in mice expressing the more potent Polγexo- variant, dsDNA segment terminated at TAS (i.e., 7S DNA) is significantly longer compared to that in the wild-type (29). Interestingly, the mice exhibit progeroid phenotype with accumulation of mtDNA deletions (79). Therefore, this excessive strand displacement ability of the Polγexo- variant could be deleterious, possibly by enabling the formation of stable secondary structures at the displaced strand that inhibit the helicase loading and promoting the strand breakage. In addition, the Polγ-mtSSB coupling would also be relevant for the processing of primers at the two origins of replication; i.e, generating long flaps upon reaching the origins of the two mtDNA strands, enabling their further processing by dedicated nucleases (80). In this context, the deleterious effect of Polγexo- could result from stimulation of mtSSB to generate excessively long flaps precluding their processing by the associated nucleases and, in turn, maturation of mtDNA.

Finally, we note that the joint destabilization energy of the first bp of the fork by the mitochondrial SSB-holoenzyme couple (∼1.4 *k*_*B*_*T* per dNTP incorporated) is comparable to that reported for replicative helicases previously, 0.05-1.6 *k*_*B*_*T* (81-83). However, the stimulation of the strand displacement activity of Polγ by mtSSB (and that of T7DNAp by gp2.5) is mainly limited by the reduced ability of the polymerase-SSB complex to counteract the fork regression pressure and therefore, control the equilibrium between the *pol* and *exo* activities efficiently. Therefore, it is tempting to speculate that during leading (or H-strand) DNA synthesis (when Polγ is assembled at the replisome) one of the main roles of the mitochondrial helicase Twinkle would be to prevent fork regression pressure on the holoenzyme, in order to ensure robust DNA replication (as proposed for other replication systems, (15,31,33)). Future experiments aimed to study the coordinated activity of these two molecular motors at the DNA fork will help to elucidate these questions.

## Supporting information

I Plaza et al_Supp Info

## SUPPLEMENTARY DATA

Supplementary Data are available

## ACKNOWLEDGEMENTS

We are grateful to members of B. Ibarra and G. Ciesielski labs for useful discussions.

