## Supplementary material for "Mechanism of strand displacement DNA synthesis by the coordinated activities of human mitochondrial DNA polymerase and SSB": I Plaza et al_Supp Info

¶ corresponding authors

**Strand displacement model**

Betterton and Jülicher proposed a theoretical framework to quantify the interaction potential of a helicase with the nucleic acid fork that has been used to assess the unwinding mechanism of several helicases (1-3). We described previously a generalization of this framework to analyse the pause-free velocity dependence on mechanical tension of DNA polymerases and quantify their base pair destabilization energies during strand displacement DNA synthesis (4). Here, we present a revision of the latter model that includes the effect of mechanical tension on the translocation step of the DNA polymerase along the DNA template. We have used this revised model, referred as strand displacement model in the main text, to determine the extent of fork destabilization by the mitochondrial polymerase holoenzyme and holoenzyme/SSB complex during strand displacement DNA synthesis.

The model considers that DNA replication occurs in discrete steps of 1 nucleotide ($\delta$ = 1 nt, (5-7)) during which the polymerase moves forward with a rate $k_{+}$. Under primer extension conditions (no dsDNA ahead of the polymerase), Polγ and Polγexo-, present maximum translocation rates $k_0$, of ~24 and 27 nt/s, respectively (Figure S1, (8)). Backwards movements, such as pyro-phosphorolysis, are highly improbable under our experimental conditions in the absence of PPi in solution (9,10), while exonuclease events of Polγ would be included within the pause kinetics and removed for the calculation of pause-free rates. Therefore, backwards translocation rates could be neglected in the computation of the overall replication rate.

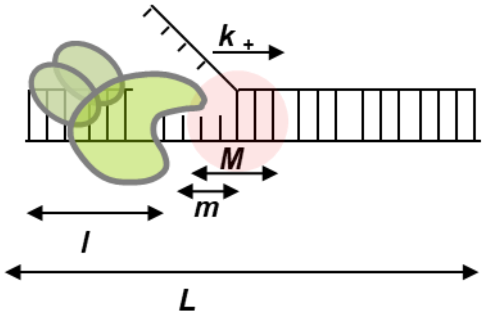


According to the strand displacement model, the pause-free strand displacement rate on a dsDNA substrate of length L is given by

$$V_{max}(\boldsymbol{l},\boldsymbol{f}) = \sum_{m=0}^{L-l} k_{+}(l,m,f)P_0(l,m,f) \text{ (Eq.1)}$$

where $P_0(l,m,f)$ is the probability of finding $m$ bases pairs open ahead of the polymerase when the enzyme is at position $l$ and the applied tension on the DNA is $f$. $P_0(l,m,f)$ is determined by the Gibbs energy, $\Delta G(l,m,f)$, required to open $m$ base pairs ahead of position $l$ as

$$P_0(l,m,f) = \frac{e^{-\Delta G(l,m,f)/k_B T}}{\sum_{m=0}^{l-l} e^{-\Delta G(l,m,f)/k_B T}} \ (Eq.2)$$

$\Delta G(l,m,f)$ is the result of three energetic contributions:

$$\Delta G(l,m,f) = \sum_{i=l+1}^{l+m} \Delta G_{bp}(i) - \Delta G_f - \min(m,M)\Delta G_{int} \ (Eq.3).$$

$\Delta G_{bp}$is the average stability of the $m$ base pairs opened ahead of the polymerase , $\Delta G_{bp}$ ~ 1.8 $k_BT$ (for our DNA hairpin) (4).

$\Delta G_f$ corresponds to the mechanical destabilization of the fork junction by mechanical tension applied on the opposite strand of the DNA hairpin. $\Delta G_f$ was defined as: $\Delta G_f = 2m\int_0^F x_{nt}(f')df'$, where $x_{nt}(f')$ is the extension of 1 ssDNA nucleotide at a given tension, and $2m$ is number of single-stranded nucleotides generated by the unwinding of $m$ base pairs of the DNA hairpin.

$\Delta G_{int}$ is the effective interaction energy of the polymerase or the polymerase/SSB complex with the dsDNA fork junction and their interaction ranges, M (red circle in the diagram above).

$k_+(l,m,f)$ is the rate of forward translocation of one nucleotide ($\delta$=1 bp) when the polymerase is at position $l$ and $m$ base pairs of the DNA hairpin are opened under mechanical tension on the DNA ($f$).

$k_+(l,m,f)$ =

$$\begin{cases} \mathbf{0}, & \boldsymbol{m < \delta} \\ \boldsymbol{k_0 \cdot exp\left[-a \cdot min(M,\delta) \cdot \frac{\Delta G_{int}}{k_B T}\right] \cdot exp\left[-\delta \cdot f \cdot \frac{\left(x_{nt}(f) - x_{bp}(f)\right)}{k_B T}\right]}, & \boldsymbol{\delta \le m < max(M,\delta)} \\ \boldsymbol{k_0 \cdot exp\left[-a \cdot (M+\delta-m) \cdot \frac{\Delta G_{int}}{k_B T}\right] \cdot exp\left[-\delta \cdot f \cdot \frac{\left(x_{nt}(f) - x_{bp}(f)\right)}{k_B T}\right]}, & \boldsymbol{max(M,\delta) \le m < M+\delta} \\ \boldsymbol{k_0 \cdot exp\left[-\delta \cdot f \cdot \left(x_{nt}(f) - x_{bp}(f)\right)/(k_B T)\right]}, & \boldsymbol{M+\delta \le m} \end{cases} \qquad (\boldsymbol{Eq.4})$$

$k_0$ corresponds to the maximum translocation rate of the holoenzyme in primer extension conditions in the absence of hairpin or secondary structures, $k_0$ ~24 and 27 nt/s for Polγ and Polγexo-, respectively (Figure S1 and (8)). The dimensionless coefficient, $a$, was previously determined by the relative location of the activation barrier for the stepping motion and was set to 0.1 (4).

The exponential term $exp\left[-\delta \cdot f \cdot \frac{\left(x_{nt}(f)-x_{bp}(f)\right)}{k_BT}\right]$ was included to account for the effect of tension on the DNA template, $f$, on the translocation rate as the polymerase translocate with a δ =1 nucleotide step size, and coverts the extension of a single-stranded nucleotide ($x_{nt}$) to that of a double-stranded ($x_{ds}$).

### Two-state model

The mitochondrial holoenzyme variants used in this study present frequent pause or non-productive states during strand displacement DNA synthesis. According to our results and previous studies (4,11), the fork regression pressure would be the main responsible for pausing transiently the advance of the polymerase, which in turn would promote the partition of the primer to the exonuclease (*exo*) site. As explained in the main text, the kinetics of the pause state(s) would be different in each holoenzyme due to the presence and absence of *exo* activity in the wild-type and *exo* deficient variants, respectively.

Idling (*pol-exo* cycles) of Polγ at the fork junction would involve few nucleotides, which cannot be resolved with our current resolution limit. Therefore, in a simplified two-state scenario, during the elongation phase of strand displacement DNA synthesis, each holoenzyme would alternate between two states; moving and pause state with an equilibrium constant, *K*. According to our results, application of increasing tension on the opposite strands of the DNA hairpin favors the moving over the pause state of the holoenzyme. Based on these results, we considered that *K* follows an Arrhenius tension dependency

$$K(f) = K(0) \cdot e^{-df} \text{ (Eq.5)},$$

where $K(0)$ is the equilibrium constant between the pause and moving states in the absence of tension (0 pN), *f* is the applied tension and *d* the conformational change along the pulling coordinate that shifts the equilibrium away from the pause state.

In terms of average residences times per nucleotide, $K(f)$ can be defined as

$$K(f) = \frac{T_p(f)}{T_a(f)} \text{ (Eq.6)}.$$

$T_a(f)$ is the average residence time in the active state, or the inverse of the tension dependent pause-free velocity, *V*(*f*), defined by Eq.1

$$T_a(f) = \frac{1}{V(f)} \text{ (Eq. 7)},$$

$T_p(f)$ is the average residence time in the pause state calculated as the difference between total average residence time per nucleotide $T(f) = \frac{1}{V_{mean}(f)}$ and $T_a(f)$. Therefore, it follows that

$$\frac{T_p(f)}{T_a(f)} = T_p(f) \cdot V(f) = K(0) \cdot e^{-df} \text{(Eq.8)},$$

and,

$$T_P(f) = \frac{K(0)e^{-df}}{V(f\ )} \text{ (Eq.9)}.$$

We note that $T_p(f)$ includes pause frequency and pause duration.

The residence time in the pause state can also be expressed in terms of moving probability (*MP*), which is the probability of finding the holoenzyme at the active state during the strand displacement reaction. *MP*(*f*) was defined as:

$$MP(f) = \frac{V_{mean(f)}}{V_{max}(f)} = \frac{T_a(f)}{T(f)} \qquad \text{(Eq. 10)}$$

and therefore,

$$MP(f) = \frac{1}{1 + K(0)e^{-df}} \qquad \text{(Eq. 11)}$$

**SUPPLEMENTARY FIGURES**

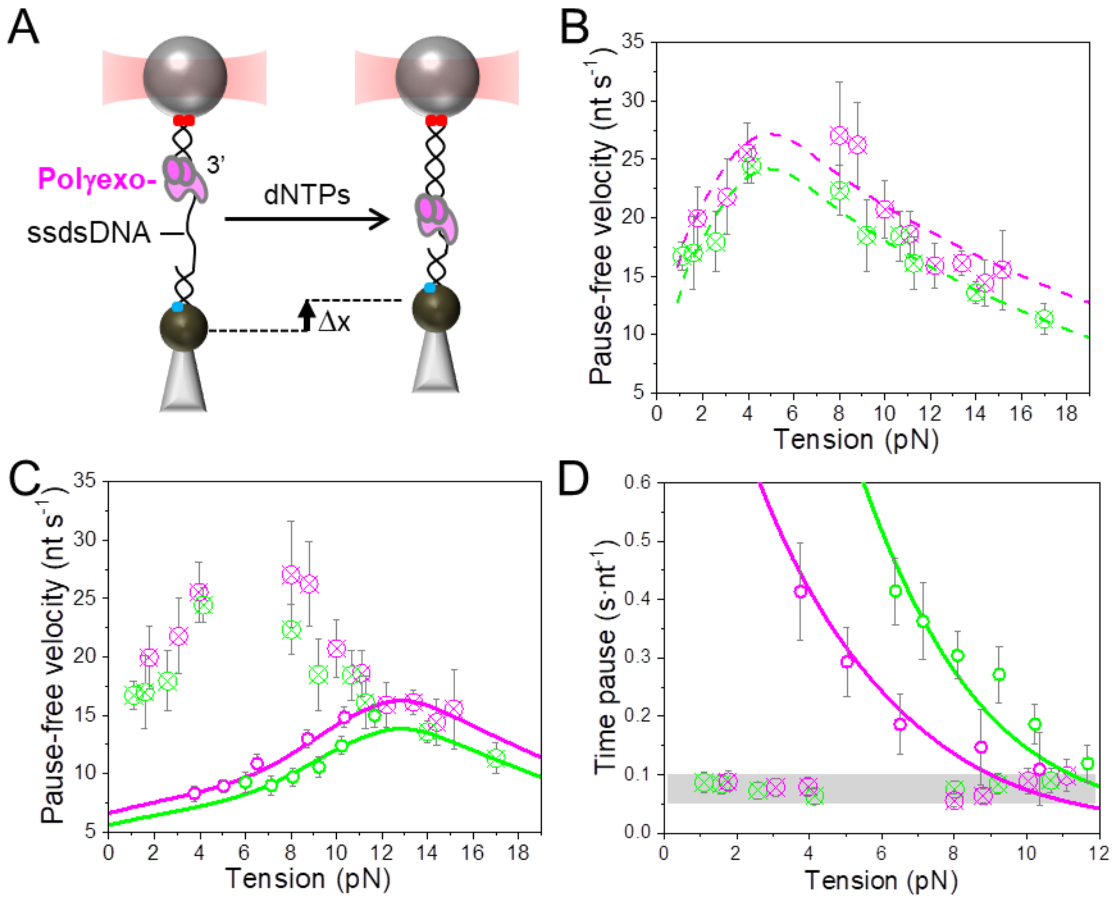


**Figure S1**: Primer extension experiments. **A)** Experimental setup to measure Polγexo- primer extension activities. A single DNA molecule containing a ssDNA gap (~900 nt) flanked by ~3550 bp dsDNA handles labelled with biotin (blue) and digoxigenin (red) moieties at their ends (described in (8)) was tethered between two functionalized plastic beads; one bead is held in the optical trap (red cone) and the other on top of a micropipette (grey). At constant tension, the activity of the polymerase changes the end-to-end extension of the DNA construct as the enzyme converts the extension of a single-stranded nucleotide to that of a double-stranded DNA (Δx). Experimental conditions were identical to those used by us to measure primer extension activities of Polγ previously (8). **B**) Average pause-free velocities of Polγ (green symbols from (8)) and Polγexo- (magenta symbols) as a function of tension (pN). Dotted lines represent fit of the data with a model considering that pause-free velocity is modulated by the work to convert a single-stranded nucleotide to double-stranded form, plus the work required to disrupt the secondary structure characteristic of free-ssDNA at each tension (8). Best fits of the model to the tension dependent pause-free rates yielded pause-free translocation rates in the absence of secondary structure of 23.0 +/- 0.7 nt/s and 27.1 +/- 12 nt/s for Polγ and Polγexo-, respectively. **C)** Comparison between Polγ (green) and Polγexo- (magenta) pause-free rates during primer extension (crossed symbols) and strand displacement conditions (circles). Solid lines represent the fit to strand displacement and primer extension data with the strand displacement model described in this work. **D)** Average residence times at pause state of Polγ (green crossed symbols) and Polγexo- (magenta crossed symbols) as a function of tension. Residences times at pause state during strand displacement conditions are shown for comparison (circles). Solid lines represent the fits with the two-state model to the residence times at pause state of Polγ (green line) and Polγexo- (magenta line) during strand displacement conditions. For clarity of display, a grey box, instead of actual values, was used in the figures of the main text to indicate the values of the average residence times at pause state during primer extension conditions. In all figures errors show s.e.

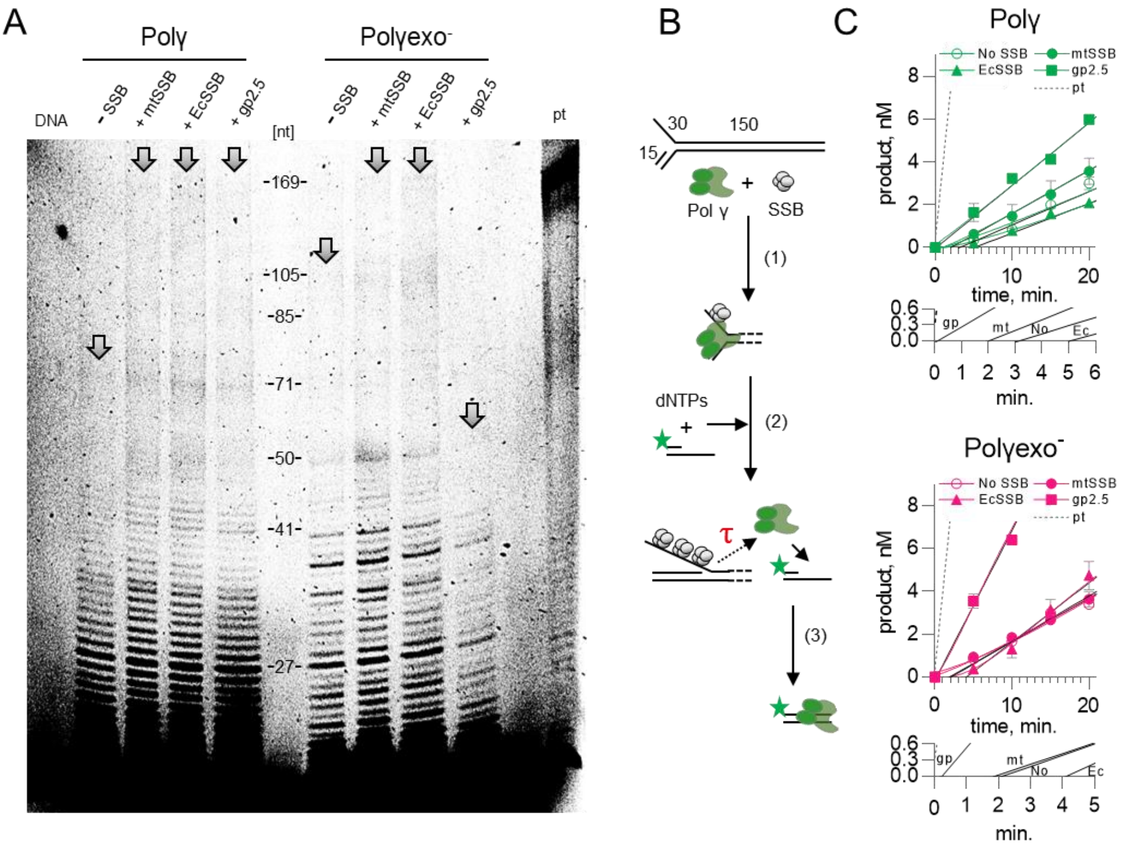


**Figure S2**: Strand displacement replication in bulk. **A)** To assess the processivity of the strand displacement replication, Polγ and Polγexo- were pre-incubated with equimolar amounts of fluorescently-labeled fork DNA substrate (depicted in B), in the presence or absence of excess SSB proteins, as indicated. DNA synthesis was initiated by the addition of dNTPs, and 100-fold excess of trap DNA (i.e. unlabeled fork DNA substrate) to limit the detection of products to those generated by a single enzyme. Reactions of Polγ and Polγexo- were stopped after 10 and 5 minutes, respectively. 'pt' lane represents the primer-template (15/169) extension (full length product) control. DNA products were analyzed on denaturing polyacrylamide gels. Arrows indicate the longest product detected in the lane. The presented image is representative of 4 repeats, using two distinct Polγ and Polγexo- preparations. **B)** Schematic representation of the strand displacement replication assay used to assess the residence time of Polγ at the fork (τ). Pol γ variants were pre-incubated with an equimolar amount of unlabeled fork DNA substrate, in the presence or absence of excess SSB proteins, enabling the assembly of the proteins at forks (1.).

DNA synthesis was initiated by the addition of dNTPs together with an excess of a fluorescently-labeled (green star) primer-template (15/44) (2.). The residence time of Polγ at the primary DNA fork substrate can be inferred from the time it takes to detect the fluorescently-labeled product, τ (3.). **C)** Assays described in B were carried out using either Polγ (green, top graph) or Polγexo- (magenta, bottom graph), in the absence (open circles) or presence of mtSSB (closed circles), E. coli SSB (closed triangles), or T7 phage gp2.5 (closed squares). Reactions were stopped at the indicated time intervals and analyzed by native polyacrylamide gel electrophoresis. The residence time of Polγ variants at the primary fork substrate/product is equivalent to the lag (τ) between the reaction initiation and the synthesis of the detectable secondary product, which was identified by fitting linear regression (black lines) to individual data sets (colored). This allowed to interpolate the X-axis intercepts, detailed in the insets below the corresponding graphs (black lines denoted by gp, mt, No, Ec), which are equivalent to the τ values. Dissociation rates ($k_{off}$) presented in Table 1. Correspond to inverted τ values. The dashed 'pt' line represents a control extension of the labeled primer-template by Polγ in the absence of the fork DNA substrate. The data represent an average of at least two experiments. Errors show s.e.

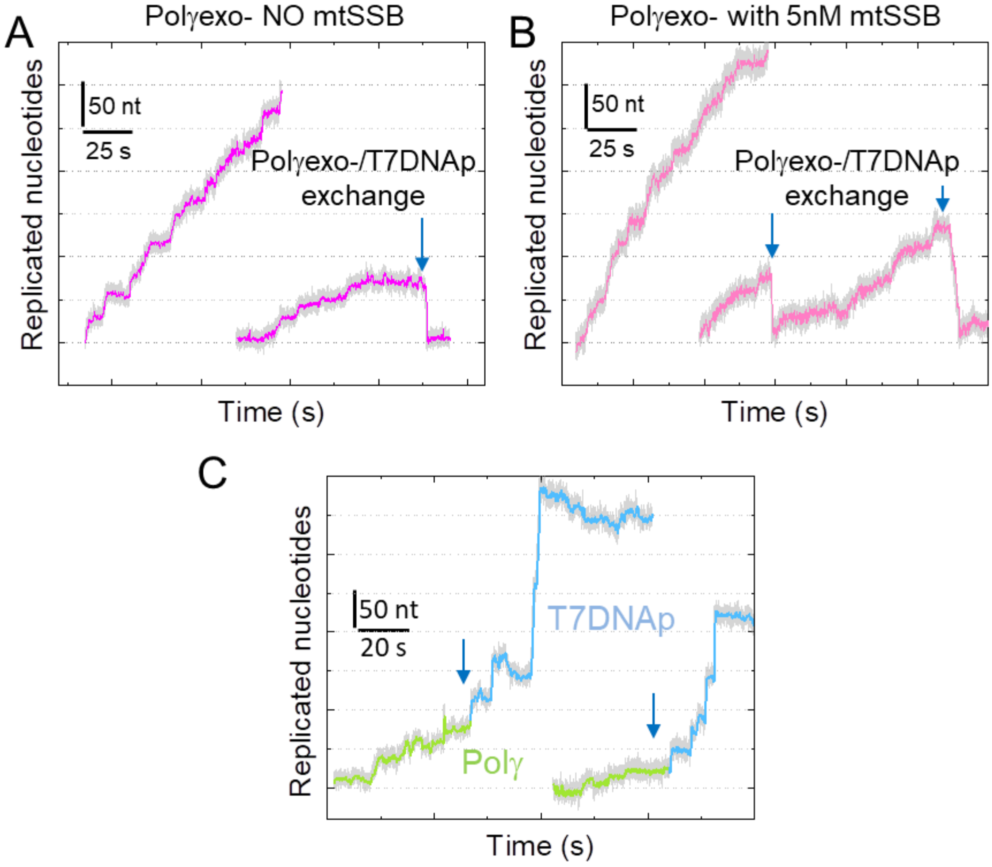


**Figure S3**: **A)** Representative strand displacement traces of the exonuclease deficient variant Polγexo- in the absence (left) and presence (right) of competing T7DNAp in solution (f~ 6 pN). **B)** Experimental traces of Polγexo- with 5 nM mtSSB in the absence (left) and presence (right) of competing T7DNAp in solution (f~ 6 pN). In **A)** and **B)** exchange of Polγexo- with T7DNAp was monitored as fast exonucleolysis events (blue arrows) not observed in the absence of T7DNAp. **C**) Representative strand displacement traces of Polγ in the presence of T7DNAp at *f*=9 pN. At tension above 8 pN, Polγ exchange by T7DNAp (blue arrow) was monitored as a sudden increase in the replication rate due to the ~4-fold difference in the replication rates of the mitochondrial (green) and phage (blue) polymerases at these tensions. For all experiments Polγexo-:T7DNAp molar ratio was 1:0.5.

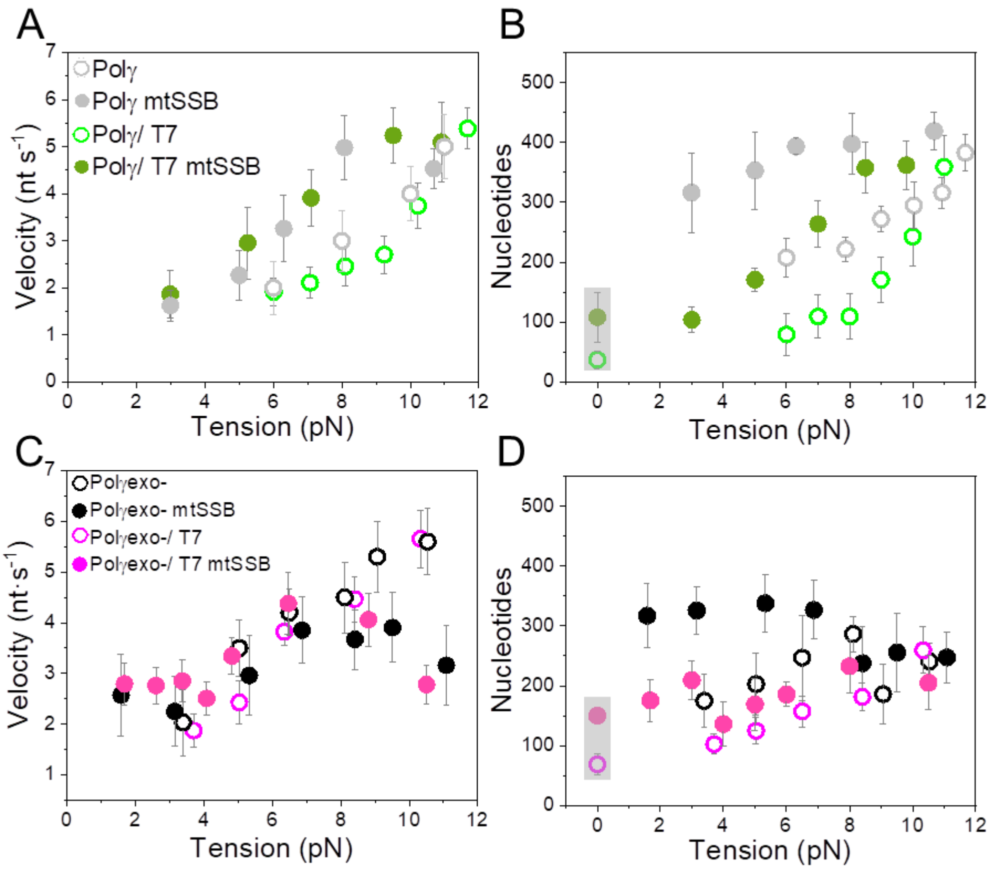


**Figure S4:** Comparisons of average number of replicated nucleotides and velocities of Polγ and Polγexo- with and without competitor T7DNAp in solution. For all figures empty dots show data in absence of mtSSB and full dots show data in the presence of 50 nM mtSSB. Note that values at 0 pN (grey boxes in B and D) correspond to single-turn over experiments in bulk. **A)** Average velocities and **B)** replicated nucleotides of Polγ as a function of tension without (grey) and with (green) of T7DNAp in solution. **C)** Average velocities and **D)** replicated nucleotides of Polγexo- as a function of tension in the absence (black) and presence (magenta) of T7DNAp in solution. In **A)** and **C)** values at 0pN correspond to single turn over bulk measurements. In all figures errors show s.e.

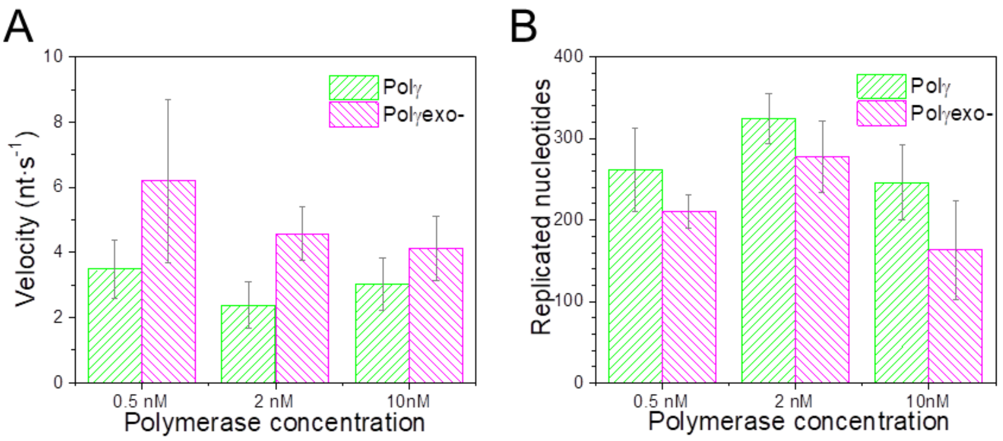


**Figure S5**: Effects of Polγ (green) or Polγexo- (magenta) concentrations on (**A**) average velocities and (**B**) number of replicated nucleotides in the absence of competitor T7DNAp in solution. Variation of holoenzyme concentration within the 0.5-10 nM range (20-fold) did not have significant effects on replication kinetics. Data was recorded at a constant mechanical tension of 8 pN. Together with the polymerase exchange experiments, these results support that under our current experimental conditions polymerase exchange is a fast, non-limiting step, which does not contribute to the observed paused kinetics significantly. Errors show s.e.

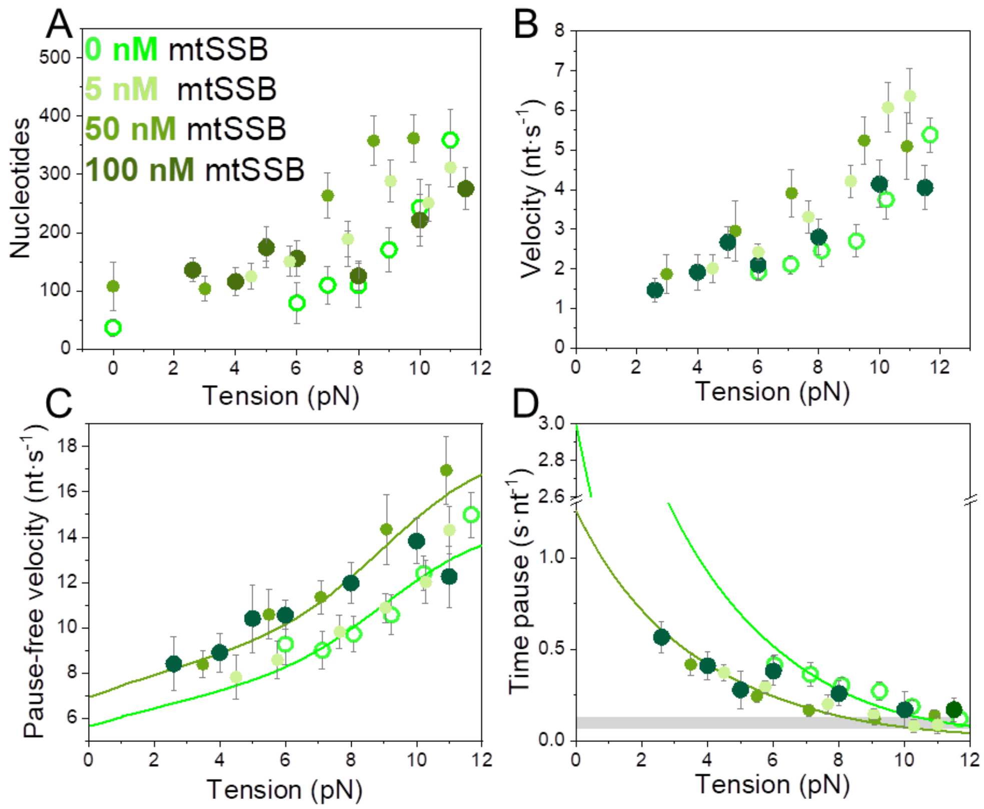


**Figure S6**: Effect of mtSSB concentration on the tension dependent strand displacement kinetics of Polγ. For all figures, empty green dots correspond to conditions with no mtSSB and dark green symbols to conditions in the presence of 100 nM mtSSB. Data in the presence of 5 nM (light green) and 50 nM (olive) mtSSB are shown as reference (more details in the main text, Figure 3). mtSSB at 100 nM concentration stimulated the average number of replicated nucleotides **(A)** and average strand displacement rates **(B)** of Polγ at tension below 6 pN. **C)** 100 nM mtSSB stimulated pause-free velocity by ~25% at tensions below 10 pN. Light and olive green lines correspond to the fits of the strand displacement model to data in the absence and presence of mtSSB (50 nM), respectively. **D)** 100 nM mtSSB decreased the average residence time at pause state per nucleotide of Polγ below ~6 pN. Light and olive green lines correspond to the fits of Eq.1 (main text) to data in the absence and presence of mtSSB (50 nM), respectively. In contrast to 50 nM mtSSB, mtSSB at 100 nM did not simulate strand displacement activity of Polγ at tension above 6 pN. For all plots, error bars show standard errors. Values at 0pN correspond to bulk measurements.

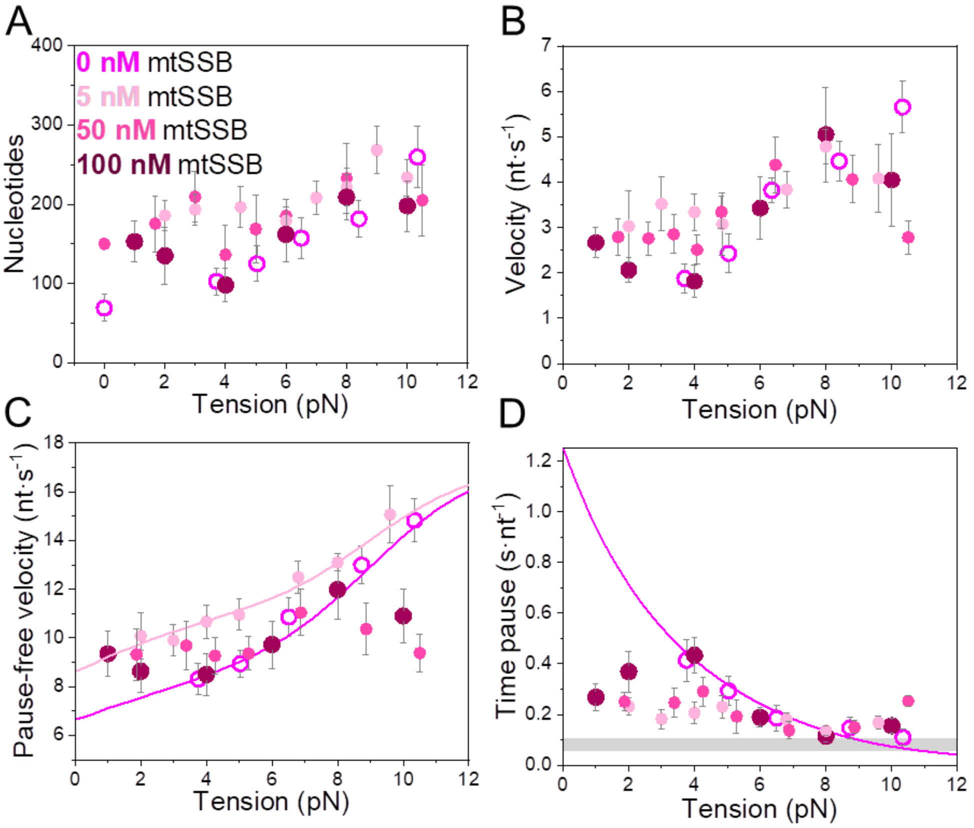


**Figure S7**: Effect of mtSSB concentration on the tension dependent strand displacement kinetics of Polγexo-. For all figures, magenta empty circles show data in the absence of mtSSB and dark purple symbols show data with 100 nM mtSSB in solution. Data in the presence of 5 nM (light pink) and 50 nM (pink) mtSSB are shown as reference and described in more detailed in Figure 4 of the main text. High mtSSB concentrations (100 nM) had a stimulatory effect on Polγexo- strand displacement activity at tension below ~4 pN. Under this tension value, mtSSB (100 nM) stimulated: **A)** the average number of nucleotides, **B)** average strand displacement rate (velocity), **C)** Pause-free velocity and **D)** decreased average residence time at pause state per nucleotide, with respect to conditions in the absence of mtSSB. However, above 4 pN, the stimulatory effects ceased and inhibitory effects were apparent above ~8pN. In **C**) magenta and light pink lines correspond to the fits of the strand displacement model to pause-free velocity data in the absence and presence of mtSSB, respectively. In **D**), magenta solid line corresponds to the fit of the two-state model (Eq.1) to the residence time at pause state per nucleotide in the absence of mtSSB. For all plots error bars show standard errors. Values at 0pN correspond to bulk measurements.

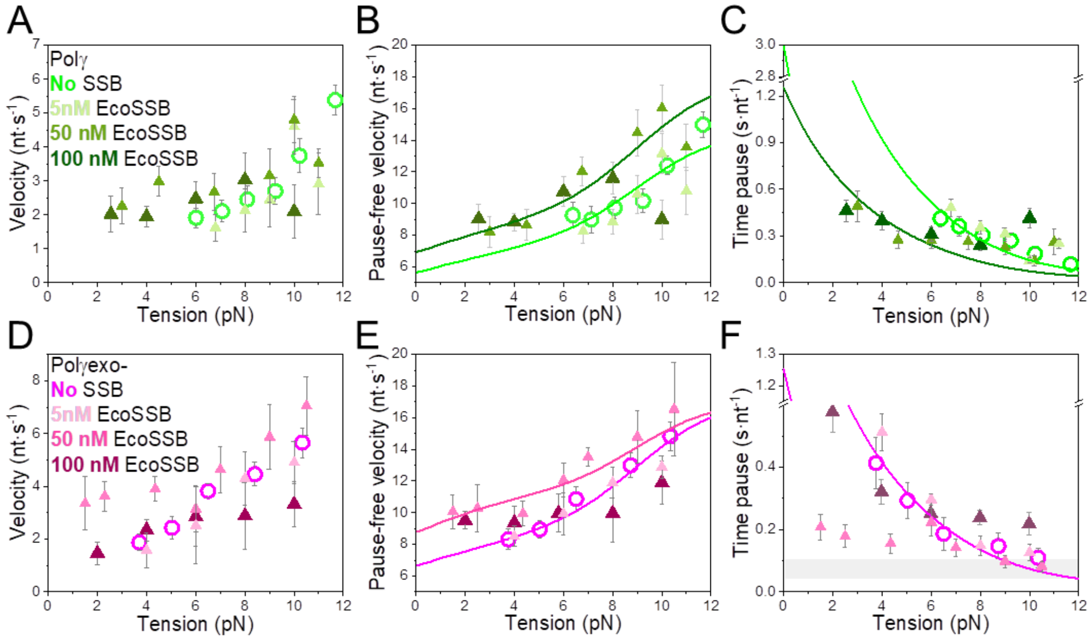


**Figure S8**: Effects of increasing concentrations of EcoSSB on Polγ and Polγexo- strand displacement kinetics. **A), B),** and **C)** effects of 100 nM EcoSSB on Polγ tension dependent strand displacement kinetics. For all figures, green empty circles show data in the absence of SSB and dark green triangles data with 100 nM EcoSSB in solution. Data in the presence of 5 or 50 nM EcoSSB (light green and olive triangles, respectively) are shown as reference and described in more detailed in Figure 5 of the main text. EcoSSB (100 nM) stimulated the average rates (**A**), pause-free velocities (**B**) and residence time at pause state (**C**) below 5-6 pN and resulted in significant inhibitory effects at high tensions (*f*>8 pN). **D), E),** and **F)** effects of 100 nM EcoSSB on Polγexo- tension dependent strand displacement kinetics. For all figures, magenta empty circles show data in the absence of SSB and dark purple triangles show data with 100 nM EcoSSB in solution. Data in the presence of 5 or 50 nM EcoSSB (light pink and pink triangles, respectively) are shown as reference (see Figure 5 of the main text for details). 100 nM EcoSSB did not change average rates (**D**), pause-free velocities (**E)** and time at the pause state (**F**) with respect to conditions in the absence of SSB, below 6 pN, and resulted in considerable inhibition at tensions above 8 pN. For all plots error bars show standard errors.

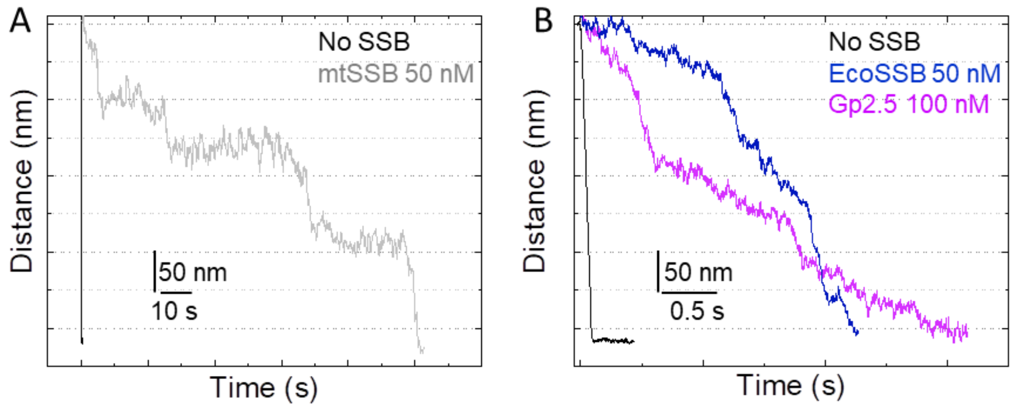


**Figure S9**: Hairpin closure times in the absence and presence of SSB. To measure the effect of SSBs on the DNA fork regression pressure, we unwound the DNA hairpin and measured its rewinding time in the absence and presence of SSBs at a constant tension of 6 pN. **A**) Representative rewinding traces in the absence (black) and presence of 50 nM mtSSB (grey). **B**) Rewinding traces in the absence (black) and presence of 50 nM EcoSSB (blue) or 100 nM gp2.5 (purple).

| | **$\Delta G_{int}$($k_B$T)/$M$** | **$K$(0)** | **$d_{T7}$(nm)** | **$T_p$(0) (s $nt^{-1}$)** |
|---|---|---|---|---|
| **T7DNAp** | 0.30 ± 0.10 /1* | 3.52 ± 1.2** | 0.84± 0.2** | 0.32** |
| **+ Gp2.5** | 0.73 ± 0.11 /1 | 2.61 ± 1.1 | 1.02± 0.3 | 0.13 |
| **+ EcoSSB** | 0.70 ± 0.10 /1 | 2.61 ± 1.1‡ | 1.02± 0.3‡ | 0.13‡ |
| **+ mtSSB** | 0.30 ± 0.10 /1* | 3.52 ± 1.2** | 0.84± 0.2** | 0.32** |
| **Sequenase©** | 0.30 ± 0.10 /1 | 3.52 ± 1.2 | 0.84± 0.2 | 0.32 |
| **+ Gp2.5** | 0.70 ± 0.11 /1 | N.A. | N.A. | ~0.09[1pN] |
| **+ EcoSSB** | 0.70 ± 0.11 /1† | N.A. | N.A. | ~0.06 [1pN] |
| **+ mtSSB** | 0.30 ± 0.10 /1* | N.A. | N.A. | 0.32 |

**Table S2**. $\Delta G_{int}/M$, minimum values of the free-parameters yielded by least squared error fits of strand displacement model to T7DNAp and Sequenase© pause-free velocity data. (*) and (†) indicated that values were obtained from fits of the strand displacement model to Sequenase© pause-free velocity data in the absence (*) or presence (†) of gp2.5. $K$(0) and $d_{T7}$ are the free-parameters of *Eq.*1, and $T_p$(0) is the average residence time at pause state in the absence of tension (0pN) obtained upon fitting experimental data in Figures 6E and 6H (main text) with the least mean squared error. (**) and (‡) values were obtained from fits of *Eq.*1 to residence time in pause state Sequenase© in the absence (**) or presence (‡) of gp2.5. ([1pN]) Average residence times at pause state at ~1 pN. In all cases, errors show standard errors. SSB concentrations were 100 nM gp2.5, 50 nM EcoSSB and 50 nM mtSSB.

<u>**Supplementary Methods**</u>

### DNA substrates for bulk experiments.

For experiments in bulk (Figure S2), DNA substrates were prepared using PAGE-purified oligonucleotides purchased from Integrated DNA Technologies or GenScript. The fork DNA substrate was prepared by combining equal concentrations of 169 nt template strand, 180 nt displaced strand containing 5'-$(dT)_{30}$, and 15 nt primer with or without 5'-Cy3 fluorescent label, in 10 mM Tris-HCl, pH 8.0, 50 mM NaCl buffer. The mixture was incubated at 95 ºC for 10 minutes and left to cool down slowly overnight. The assembled fork construct contains a 150 bp duplex segment separated from the upstream 15 nt primer by a 4 nt gap and preceded by a 5'-$(dT)_{30}$ overhang of the displaced strand. The assembly and quality of the fork DNA substrate was confirmed by visualization on 10% native polyacrylamide gel. For the fork residence time analysis (Figure S2C), the secondary primer-template substrate was prepared by annealing the 5'-Cy3-labeled 15 nt primer to a 44 nt oligonucleotide as described above.

| | |
|---|---|
| **169 nt template oligomer** (IDT) | ATTAGACTGAACACGAACTGGATGCTACCTGAAGTGATTGATTACG ATGAACGTGAACTGGATGCTACCTATTAGACTGAACACGAACTGGA TGCTACCTGAAGTGATTGATTACGATGAACATGAACTCGATGCTAC CTGAAGTGATTGGACTGGGAAAACCCTGGCG |
| **180 nt displaced oligomer** (IDT) | TTTTTTTTTTTTTTTTTTTTTTTTTTTTTTCAATCACTTCAGGTAGCATC GAGTTCATGTTCATCGTAATCAATCACTTCAGGTAGCATCCAGTTC GTGTTCAGTCTAATAGGTAGCATCCAGTTCACGTTCATCGTAATCAA TCACTTCAGGTAGCATCCAGTTCGTGTTCAGTCTAAT |
| **15 nt primer +/- 5'-Cy3** (GenScript) | CGCCAGGGTTTTCCC |
| **44 nt template oligomer** (IDT) | GCACTGGCCGTCGTTTTACGGTCGTGACTGGGAAAACCCTGGCG |

**Table S1:** Sequences given in 5'-3' direction.

### Bulk biochemical assays

*Strand displacement replication processivity assay*. Reactions were performed in 50 mM Tris HCl, pH 8.5, 30 mM KCl, 10 mM DTT, 4 mM $MgCl_2$, 0.1 mg/mL bovine serum albumin, and 10% glycerol. In a single assay, 50 nM wild-type or exo- Polγ was preincubated with 50 nM Cy3-labeled fork DNA substrate, in the presence or absence of 750 nM mtSSB, 250 nM *Eco*SSB, or 1 µM gp2.5 for 3 minutes at room temperature. Concentrations of the SSB proteins correspond to, respectively, three-, one-, and two-fold excess of assembled multimers over the concentration required to cover the displaced strand of the fork substrate fully. Also, stoichiometries of mtSSB and EcoSSB proteins to DNA substrate correspond to peak stimulation of Polγ activity, reported before (12). Samples were next moved to 37 ºC and DNA synthesis was initiated by the addition of dNTPs to the final concentration of 400 µM, and excess DNA trap (1.5 µM unlabeled fork DNA substrate) to prevent the exchange of polymerases at the originally-bound fork (approach similar to the one applied in (13)). Reactions with Polγ and $Polγexo^{-}$ were stopped after 10 and 5 minutes, respectively, by the addition of 10x stop solution (80 mM EDTA, 0.08% SDS) and 0.5 µg of Proteinase K (ThermoFisher Scientific), followed by 20 minutes incubation at 55 ºC and addition of equal volume of 90% formaldehyde, 50 mM EDTA solution. DNA was denatured at 95°C for 10 min. and analyzed on 12% denaturing (7M

Urea) polyacrylamide (19:1) gels. The gels were scanned for Cy3 fluorescence in a ChemiDoc MP Imager (BioRad). The length of products above 50 nt was interpolated from an exponential decay trend fitted to the distribution of bands corresponding to ≤50 nt, and to the 75 and 200 nt bands of a DNA marker ran in parallel on the same gel and visualized by ethidium bromide staining after the fluorescence scanning. The maximal processivities were calculated by subtracting the length of the primer (15 nt) and the gap (4 nt) from the length of the longest specie detected.

*Time course assay for the residence time of Polγ at the DNA fork*. Reactions were performed in 50 mM Tris HCl, pH 8.5, 30 mM KCl, 10 mM DTT, 4 mM $MgCl_2$, 0.1 mg/mL bovine serum albumin, and 10% glycerol. In a single assay, 10 nM wild-type or exo- Pol γ holoenzyme was pre-incubated with 10 nM unlabeled fork DNA substrate, in the presence or absence of 150 nM mtSSB, 50 nM *Eco*SSB, or 400 nM gp2.5, for 3 minutes at room temperature. SSB stoichiometries correspond to, respectively, three-, one-, and three-fold excess of assembled multimers over the concentration required to cover the displaced strand of the fork substrate fully. Samples were then moved to 37 ºC and DNA synthesis was initiated by the addition of dNTPs to the final concentration of 100 µM, and 50 nM Cy3-labeled 15/44 nt primer-template DNA (see the diagram in Figure 2B). For the control, primer-template extension in the absence of the primary fork substrate was also assessed (referred to as 'pt' in the corresponding figure). The reactions were stopped at the indicated time intervals, by the addition of 10x stop solution (100 mM EDTA, 6% SDS, 50% glycerol) and 0.5 µg of Proteinase K (ThermoFisher Scientific), followed by further incubation at 37 ºC for 30 minutes. Samples were next analyzed on native 12 % polyacrylamide (29:1) gel. Gels were scanned for Cy3 fluorescence in a ChemiDoc MP Imager (BioRad). The relative abundance of the double stranded product at each time point was measured by Image J (NIH) and plotted in GraphPad Prism. The asymmetric sigmoidal curve (presented in Figure S2C as the colored trend line) was then fitted to the full data range (0-100 min), followed by fitting a linear regression trend to the linear phase of the asymmetric curve (i.e., from 600 s for Polγ and from 300 s for Polγex0- data, with the exception of the plot for the Polγexo- with gp2.5 and the pt control, in which cases the line was fitted to all initial values). The x-axis intercept points were interpolated from the linear trend line. These values are equivalent to the lag time ($\tau$) preceding the synthesis of the detectable secondary product at constant rate, or, conversely, to the time of residence of Polγ at the primary undetectable fork DNA substrate. The dissociation rates ($k_{off}$) presented in Table 1 correspond to inverted $\tau$ values.
